# Dissecting HSV-1-induced host shut-off at RNA level

**DOI:** 10.1101/2020.05.20.106039

**Authors:** Caroline C. Friedel, Adam W. Whisnant, Lara Djakovic, Andrzej J. Rutkowski, Marie-Sophie Friedl, Michael Kluge, James C. Williamson, Somesh Sai, Ramon Oliveira Vidal, Sascha Sauer, Thomas Hennig, Bhupesh Prusty, Paul J. Lehner, Nicholas J. Matheson, Florian Erhard, Lars Dölken

## Abstract

Herpes simplex virus 1 (HSV-1) installs a profound host shut-off during lytic infection. The virion host shut-off (*vhs*) protein plays a key role in this process by efficiently cleaving both host and viral mRNAs in a translation-initiation-dependent manner. Furthermore, the onset of viral DNA replication is accompanied by a rapid decline in transcriptional activity of the host genome. Both mechanisms have tremendous impact on the RNA expression profile of the infected cells. To dissect their relative contributions and elucidate gene-specific host transcriptional responses throughout the first 8h of lytic HSV-1 infection, we here employed RNA-seq of total, newly transcribed (4sU-labelled) and chromatin-associated RNA in wild-type (WT) and Δ*vhs* infection of primary human fibroblasts. Following virus entry, v*hs* activity rapidly plateaued at an elimination rate of around 30% of cellular mRNAs per hour until 8h p.i. In parallel, host transcriptional activity dropped down to 10-20%. While the combined effects of both phenomena dominated infection-induced changes in total RNA, extensive gene-specific transcriptional regulation was observable in chromatin-associated RNA. This was surprisingly concordant between WT HSV-1 and its Δ*vhs* mutant and at least in parts mediated by the embryonic transcription factor DUX4. Furthermore, both WT and Δ*vhs* infection induced strong transcriptional up-regulation of a small subset of genes. Most of these were either poorly or not at all expressed prior to infection but already primed by H3K4me3 histone marks at their promoters. Most interestingly, analysis of chromatin-associated RNA revealed *vhs*-nuclease-activity-dependent transcriptional down-regulation of at least 150 cellular genes, in particular of many genes encoding integrin adhesome and extracellular matrix components. This was accompanied by a *vhs*-dependent reduction in protein levels by 8h p.i. for many of these genes. In summary, our study provides a comprehensive picture of the molecular mechanisms that govern cellular RNA metabolism during the first 8h of lytic HSV-1 infection.

**Author Summary:** The HSV-1 virion host shut-off (*vhs*) protein efficiently cleaves both host and viral mRNAs in a translation-dependent manner. In this study, we model and quantify changes in *vhs* activity as well as the virus-induced global loss of host transcriptional activity during productive HSV-1 infection. In general, HSV-1-induced alterations in total RNA levels were found to be predominantly shaped by these two global processes rather than gene-specific regulation. In contrast, chromatin-associated RNA depicted gene-specific transcriptional changes. This revealed highly concordant transcriptional changes in WT and *Δvhs* infection, confirmed DUX4 as a key transcriptional regulator in HSV-1 infection and depicted *vhs*-dependent, transcriptional down-regulation of the integrin adhesome and extracellular matrix. The latter explained some of the gene-specific effects previously attributed to *vhs*-mediated mRNA degradation and resulted in a concordant loss in protein levels by 8h p.i. for many of the respective genes.

## Introduction

Herpes simplex virus 1 (HSV-1), one of eight herpesviruses infecting humans, is widely known for causing cold sores but also associated with life-threatening diseases, such as encephalitis [1, 2]. A key characteristic of HSV-1 lytic infection is the induction of a profound host shut-off that is predominantly installed at the RNA level. The virion host shut-off (*vhs*) endonuclease plays a crucial role in this process. *Vhs* is delivered by the tegument of the incoming virus particles and, together with *de novo* expressed *vhs* protein, rapidly starts cleaving both cellular and viral mRNAs in a translation-initiation-dependent manner [3–8]. Later on in infection, *vhs* nuclease activity is dampened by the concerted action of at least two viral proteins, i.e. UL48 (VP16) and UL49 (VP22) [9–11], with the viral UL47 protein (VP13/14) potentially also being involved [12]. In addition to *vhs*-mediated mRNA degradation, HSV-1 shuts down host gene expression by efficiently recruiting RNA polymerase II (Pol II) and elongation factors from the host chromatin to the replicating viral genomes [13–15]. This results in an extensive loss of Pol II occupancy from host chromatin starting with the advent of viral DNA replication by 2-3h post infection (h p.i.). Furthermore, HSV-1 induces proteasome-dependent degradation of Pol II later on (>12h p.i.) in infection [16]. Finally, extensive RNA degradation upon cleavage by the *vhs* nuclease also appears to contribute to the transcriptional shut-off by 24h of infection [17].

Both *vhs*-mediated mRNA degradation and global inhibition of transcription substantially alter the host transcriptome during productive infection. Virus-induced alterations in total RNA levels can be a consequence of either of these two global phenomena or due to gene-specific changes in RNA stability or transcription, but the relative contribution of each could so far not be distinguished. Recently, Pheasant *et al*. presented a genome-scale RNA-seq study analyzing nuclear-cytoplasmic compartmentalization of viral and cellular transcripts during lytic HSV-1 infection [18]. They proposed that the translational shut-off induced by HSV-1 is primarily a result of *vhs*-induced nuclear retention and not degradation of infected cell mRNA. Furthermore, they suggested differential susceptibility of transcripts to *vhs* RNA cleavage activity. We previously performed 4-thiouridine (4sU) labeling followed by sequencing (4sU-seq) to characterize *de novo* transcription and RNA processing in hourly intervals during the first 8h of lytic HSV-1 infection of primary human foreskin fibroblasts (HFF) (Fig 1A) [19, 20]. This revealed extensive transcription downstream of genes resulting from disruption of transcription termination (DoTT) for the majority of but not all cellular genes. Due to nuclear retention of the respective aberrant transcripts, DoTT also notably contributes to host shut-off [20]. Furthermore, read-in transcription from upstream genes commonly results in the seeming induction of genes. DoTT and read-in transcription thus confounds the analysis of changes in host transcriptional activity during HSV-1 infection.

**Figure 1:**
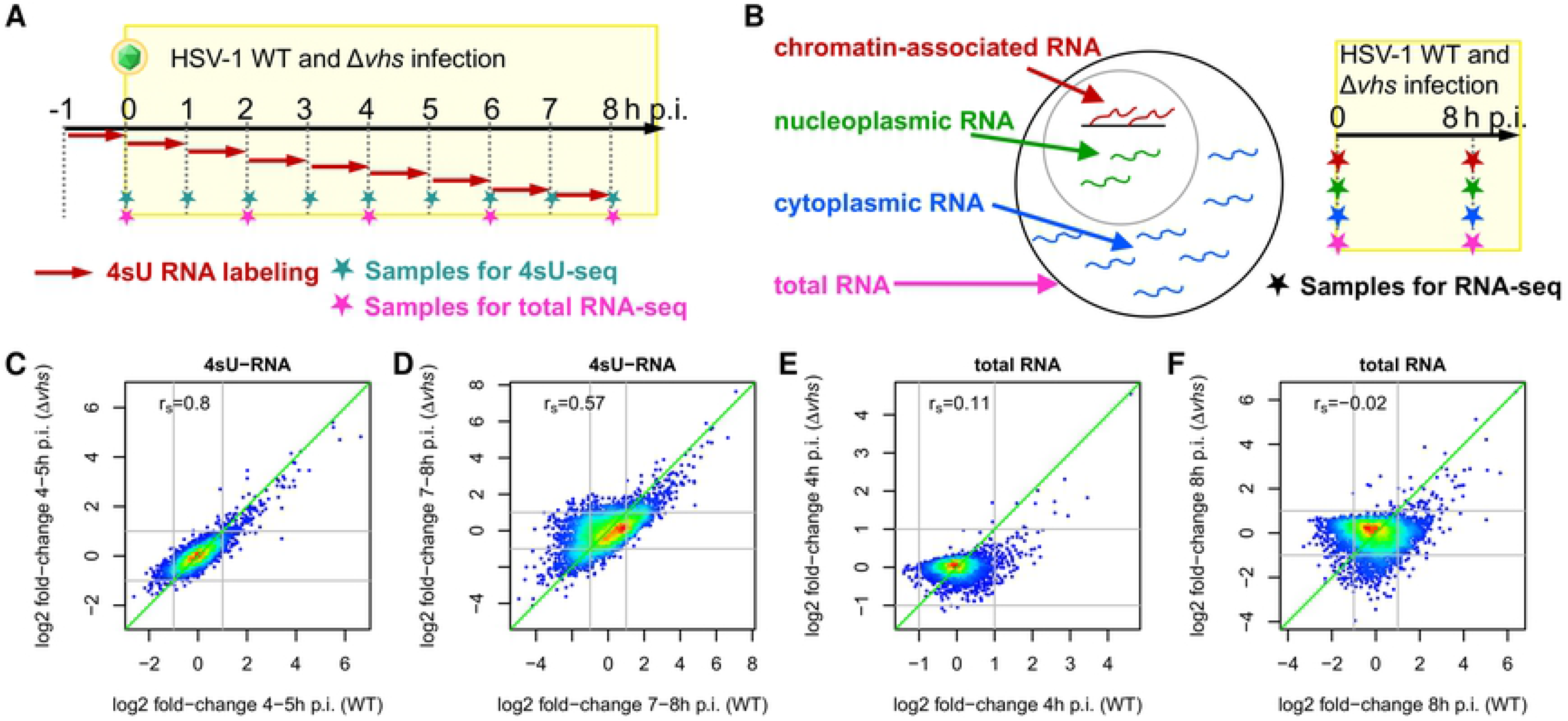
Experimental set-up and correlation of gene expression changes. (A-B) Experimental set-up of the 4sU-seq and total RNA time-courses (A) and sequencing of subcellular RNA fractions (B) in HSV-1 WT and *Δvhs* infection. The time-course experiments of the two viruses were performed as two independent experiments. Infections for the subcellular RNA fractions were performed within the same experiment. Data for WT infection for both experiments have already been published [19, 20]. (C-F) Scatterplots comparing log2 fold-changes in gene expression (infected vs. mock) between WT infection (x-axis) and *Δvhs* infection (y-axis) for 4sU-seq RNA from 4-5h p.i. (C) and 7-8h p.i. (D) as well as for total RNA from 4h p.i. (E) and 8h p.i. (F). Points are color-coded according to density of points: from red = high density to blue = low density. Spearman rank correlation *r*_*s*_ is shown on the top-left of each panel.

To dissect the effects of *vhs*-mediated RNA degradation and global loss in transcriptional activity during lytic HSV-1 infection on a genome-wide scale, we now performed total RNA-seq and 4sU-seq time-course analysis on HFF infected with a *vhs*-null mutant virus (Δ*vhs*) using the exact same experimental setting as previously employed for wild-type (WT) HSV-1 infection (Fig 1A) [19]. Furthermore, we analyzed subcellular RNA fractions (cytoplasmic, nucleoplasmic and chromatin-associated RNA) at 0 and 8h p.i. of WT and Δ*vhs* infection (Fig 1B). Mathematical modelling of RNA synthesis and *vhs*-mediated RNA decay revealed that *vhs* activity rapidly plateaued upon WT HSV-1 infection with *vhs* continuously degrading about 30% of cellular mRNAs per hour until at least 8h p.i. In contrast, total RNA changes in Δ*vhs* infection were dominated by the global loss in Pol II activity. Changes in total mRNA levels upon HSV-1 infection are thus shaped by differences in basal transcription and RNA turnover rates between the individual genes. In contrast, chromatin-associated RNA provided an unbiased picture of gene-specific transcriptional changes. This revealed an extensive, previously unsuspected *vhs*-dependent transcriptional down-regulation of the integrin adhesome and extracellular matrix (ECM). Notably, this included the key *vhs*-sensitive genes reported by Pheasant *et al*. Accordingly, increased reduction of total mRNA levels for these genes is not due to increased susceptibility to *vhs*-mediated RNA decay of the respective transcripts, but rather due to additional, *vhs*-cleavage-activity-dependent effects on their transcription. Strikingly, *vhs*-dependent down-regulation of transcriptional activity resulted in reduced protein levels of many of the respective genes already at 8h p.i. in WT but not in *Δvhs* infection as confirmed by quantitative whole-proteome mass spectrometry.

## Results

### Genome-wide RNA-seq analysis in WT and Δ*vhs* infection

To dissect the role of *vhs*, global inhibition of Pol II activity and host gene-specific regulation during productive HSV-1 infection, we employed the same experimental set-up for Δ*vhs* virus as for our previous transcriptome analyses on WT HSV-1 infection [19]. We infected HFF with Δ*vhs* at a high MOI of 10 and performed 4sU-seq in hourly intervals and total RNA-seq every two hours during the first 8h of infection (2 biological replicates; Fig 1A). It is important to note that although the WT and Δ*vhs* time-course experiments were performed independently, we carefully standardized the experimental conditions, e.g. by infecting the same batch of cells following the same number of splits after thawing as well as using the same batch of fetal bovine serum (FBS), to achieve a maximum level of reproducibility. Consistent with our previous findings [19, 21] and with the modest attenuation of Δ*vhs* in HFF, HSV-1-induced DoTT and subsequent poly(A) read-through transcription in Δ*vhs* infection was similar but slightly reduced compared to WT infection (Fig A,B in S2 File, S3 Dataset). Since read-in transcription into downstream genes due to HSV-1-induced DoTT from upstream genes can be mistaken for “induction” of these downstream genes [19], we excluded genes with read-in transcription from all following analyses (see methods for details). This resulted in a set of 4,162 genes without read-in transcription for which RNA fold-changes comparing infection vs. mock and their significance were determined using DESeq2 [22]. It should be noted that these RNA-seq fold-changes only indicate relative changes in RNA abundance compared to other genes and do not depict global reductions in RNA levels that affect all genes equally.

### Delineating *vhs*-mediated RNA degradation and loss of transcriptional activity

Gene expression fold-changes in 4sU-RNA were highly correlated between *Δvhs* and WT infection when comparing the same time points, confirming the high degree of standardization between the two independent experiments (Fig 1C,D, Fig C in S2 File). The only exceptions were the first two 4sU-seq time points (0-1 and 1-2h p.i.), when essentially no (n≤2) cellular genes were differentially expressed in both WT and *Δvhs* infection (multiple testing adjusted p≤0.001, |log2 fold-change| ≥1). This was expected as fold-changes were only very small (median |log2 fold-change| ≤0.1) and dominated by experimental noise. The highest correlations between 4sU-seq fold-changes in WT and *Δvhs* infection compared to mock were observed at 4-5h and 5-6h p.i. (Spearman rank correlation *r*_*s*_ ≈ 0.8, Fig 1C, Fig C in S2 File). Correlations decreased towards the end of the time-course in particular for genes down-regulated in WT (Fig 1D), consistent with the well described effects of *vhs* on cellular RNA levels late in infection [23]. Notably, the later stages of Δ*vhs* infection (from 6-7h p.i.) were better correlated to slightly earlier stages (4-5h, 5-6h p.i.) of WT infection (Fig C in S2 File), indicating slightly slower progression of *Δvhs* infection.

In contrast to 4sU-RNA, fold-changes in total RNA obtained from WT and Δ*vhs* infection were only poorly correlated (*r*_*s*_ ≤ 0.11, Fig 1E,F, Fig D in S2 File). Consistent with the cleavage activity of *vhs*, this was particularly prominent for genes down-regulated in WT infection. As 4sU-RNA was purified from total RNA, the poor correlation for total RNA fold-changes cannot be explained by poor reproducibility between the two independent experiments. We conclude that this instead reflects the expected strong impact of *vhs* cleavage activity on the cellular mRNAs. In principle, *vhs* cleavage activity should more strongly affect total mRNA levels of long-lived mRNAs than of short-lived mRNAs, as the former have much weaker *de novo* transcription relative to total RNA levels and are thus much more slowly replaced. On the contrary, HSV-1-induced global loss in transcriptional activity should more strongly affect total RNA levels of unstable, short-lived mRNAs. To test this hypothesis, we correlated the observed changes in total RNA upon WT and Δ*vhs* infection with the RNA half-life of the respective transcripts. RNA half-lives were obtained based on newly transcribed RNA to total RNA ratios from uninfected HFF as previously described [24]. This revealed the expected striking differences between WT and *Δvhs* infection. In WT infection, total RNA fold-changes and mRNA half-lives were negatively correlated (*r*_*s*_ = −0.38 at 8h p.i., Fig 2A), i.e. total RNA levels of stable cellular mRNAs tended to decrease more strongly than of unstable mRNAs. This was already observable at 2h p.i. (*r*_*s*_ = −0.31) consistent with mRNA cleavage and degradation by tegument-delivered *vhs* protein. The negative correlation to RNA half-lives was also confirmed for total RNA fold-changes from the study of Pheasant *et al*. at 4h p.i. (*r*_*s*_ = −0.36, Fig E in S2 File), while at 12h p.i., a weaker, but still highly significant, negative correlation was observed (*r*_*s*_ = −0.15).

**Figure 2:**
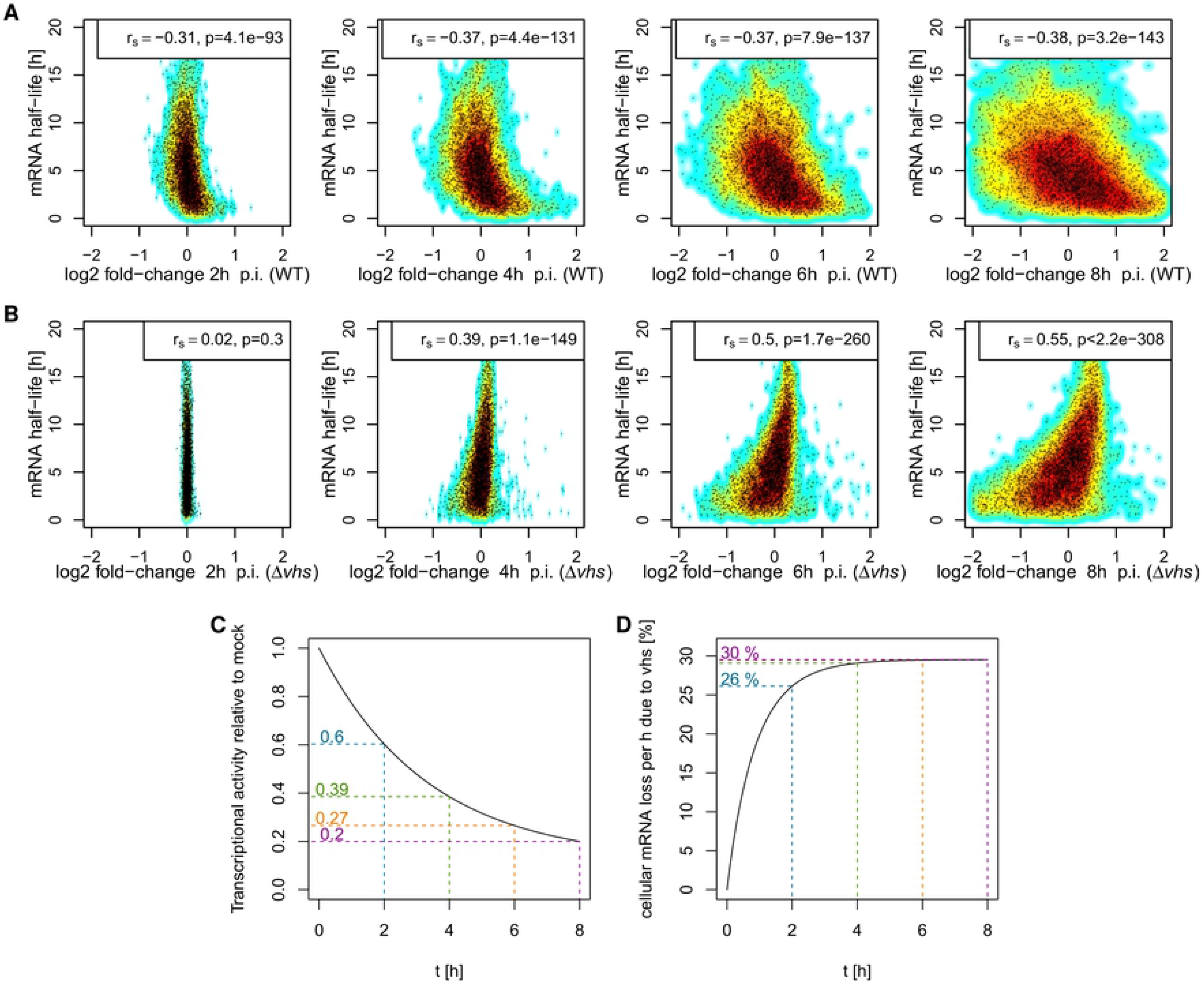
Effects of *vhs* activity and loss of transcriptional activity. (A-B) Scatterplots comparing log2 fold-changes in total RNA at 2, 4, 6 and 8h p.i. (x-axis), respectively, against RNA half-lives (y-axis) for WT (A) and *Δvhs* (B) infection. Background indicates density of points: from dark red=high density to cyan=low density. Spearman rank correlation *r*_*s*_ and p-value for significance of correlation is shown on the top of each panel. (C) Decrease in transcriptional activity relative to uninfected cells (y-axis) during HSV-1 infection (x-axis=h p.i.) estimated with our mathematical model from total RNA-seq data in *Δvhs* infection (see S1 Text). (D) Development of *vhs* activity over time as estimated with our mathematical model from total RNA-seq data in WT infection (assuming the same decrease in transcriptional activity as for *Δvhs* infection, see S1 Text). x-axis indicates h p.i. and y-axis shows the rate of cellular mRNA loss per hour (in %) due to *vhs* activity.

In *Δvhs* infection, total RNA fold-changes and RNA half-lives were positively correlated from 4h p.i. onwards (*r*_*s*_ = 0.55 at 8h p.i., Fig 2B). Thus, total RNA levels of short-lived cellular RNAs were more strongly reduced than of long-lived ones. This effect is consistent with the well described gradual decline in global transcriptional activity starting around 3-4h p.i. [15, 19]. Accordingly, total RNA fold-changes in Δ*vhs* infection largely reflect the global loss in transcriptional activity during lytic HSV-1 infection rather than gene-specific regulation. The presence of negative correlations in WT infection, however, suggests that *vhs*-mediated RNA decay, not the global reduction in transcriptional activity on cellular genes, dominates total RNA fold-changes in lytic WT HSV-1 infection. Interestingly, negative correlations in WT infection and positive correlations in Δ*vhs* infection to RNA half-lives were also observed for total RNA fold-changes between 2 and 4h p.i., 4 and 6h p.i. and 6 and 8h p.i. (Fig F in S2 File). This indicates that *vhs* cleavage activity is not as rapidly silenced upon initiation of viral gene expression as previously thought, but continues to dominate changes in total RNA levels at least until 8h p.i. Nevertheless, the much weaker negative correlation at 12h p.i. observable in the data of Pheasant *et al*. are consistent with a near complete loss of *vhs-*mediated cleavage activity at later times of infection by the combined action of the viral VP16 and VP22 proteins [9–11].

To quantify *vhs* activity throughout infection, we developed a mathematical model to estimate both the extent of loss in transcriptional activity as well as *vhs* endonuclease activity during HSV-1 infection (S1 Text) based on our total RNA-seq time-course data for 0, 2, 4, 6 and 8h p.i. in WT and *Δvhs* infection. Our results indicate that by 8h p.i., transcriptional activity dropped down to 10-20% of the level in uninfected cells during *Δvhs* infection (Fig 2C). Assuming an at least similar drop in transcriptional activity in WT infection, our model suggests that at the height of *vhs* activity, ~30% of RNAs are lost per hour due to *vhs*-mediated degradation (Fig 2D). This rate reached 26% as early as 2h p.i. and remained fairly constant until 8h p.i. Our data exclude a significant drop in *vhs* activity before 8h p.i. as the drop in transcriptional activity would otherwise result in positive correlations between total RNA fold-changes and mRNA half-lives in WT infection (S1 Text). Similarly, if the loss of transcriptional activity in WT infection were dramatically higher than in *Δvhs* infection, *vhs*-mediated degradation would have to increase even faster and to higher levels to achieve the observed negative correlations.

Although statistically significant correlations were also observed between 4sU-RNA fold-changes and RNA half-lives, these were relatively small (Fig G in S2 File) in both WT (*r*_*s*_ ≥ −0.15) and *Δvhs* infection (*r*_*s*_ ≤ 0.25). Thus, changes in newly transcribed RNA obtained during 60min of 4sU-labeling are also influenced by *vhs-*mediated decay and loss of transcriptional activity, but substantially less strongly than for total RNA. In summary, these results indicate that the poor correlation in total RNA fold-changes between WT and *Δvhs* infection is a direct consequence of global effects of *vhs* on RNA stability throughout the first 8h of lytic infection. Accordingly, our model implies that the wide range of total RNA fold-changes observed between genes in HSV-1 infection can be largely explained by differences in RNA half-lives between genes and does not require extensive gene-specific differences in *vhs-*mediated mRNA cleavage.

### Chromatin-associated RNA allows unbiased quantification of transcriptional regulation during HSV-1 infection

Since our analysis showed some effect of *vhs-*mediated decay and loss of transcriptional activity on 4sU-RNA, we analyzed subcellular RNA fractions (cytoplasmic, nucleoplasmic and chromatin-associated RNA) from mock-, WT- and Δ*vhs*-infected cells at 8h p.i. (n=2; Fig 1B) to obtain an unbiased picture of transcriptional activity in WT and Δ*vhs* infection. Here, subcellular fractions for mock-, WT- and Δ*vhs*-infected cells were obtained and sequenced in the same experiment. Only the data from mock and WT-infected cells have previously been published [20]. The efficient separation of the cytoplasmic from the nuclear RNA fraction was confirmed by the enrichment of well-described nuclear lincRNAs (MALAT1, NEAT1, MEG3) in nucleoplasmic and chromatin-associated RNA as well as cytoplasmic enrichment of reported cytoplasmic lincRNAs (NORAD, VTRNA2-1; Fig H in S2 File). In addition, overrepresentation of intronic reads in chromatin-associated RNA compared to nucleoplasmic RNA confirmed the efficient separation of these two RNA fractions (Fig H in S2 File). The substantial increase in intronic reads in the nucleoplasmic RNA fraction in WT infection is due to extensive poly(A) read-through, which results in read-in transcription into downstream genes, incomplete splicing of read-through transcripts and nuclear retention of read-through transcripts [19, 20]. This was also observed in *Δvhs* infection, however less pronounced. Notably, analysis of total RNA sequenced in addition to subcellular fractions in this experiment confirmed the results from our time-course experiments (Fig I in S2 File). Thus, the poor correlation of total RNA fold-changes between WT and Δ*vhs* infection does not result from experimental bias between two independently performed experiments. Furthermore, negative and positive correlations to RNA half-lives were again observed for WT and Δ*vhs* infection, respectively.

Since chromatin-associated RNA remains attached to the chromatin by the actively transcribing polymerases, it is not accessible to *vhs*-mediated RNA cleavage and degradation. Furthermore, it represents nascent RNA synthesized in a very short time interval in which only a negligible change in global transcriptional activity occurs. This is evidenced by the absence of any significant correlation between fold-changes in chromatin-associated RNA and RNA half-life for both WT and Δ*vhs* infection (*r*_*s*_ =−0.08 for WT and 0.07 for *Δvhs* infection), which provides further evidence for the efficient separation of the chromatin-associated RNA fraction. We thus focused on changes in chromatin-associated RNA to assess the effects of HSV-1 infection and *vhs* on transcriptional regulation. Strikingly, comparison of chromatin-associated RNA fold-changes revealed that changes in gene-specific transcriptional activity at 8h p.i. were extremely similar between WT and *Δvhs* infection (*r*_*s*_ = 0.89, Fig 3A). Thus, although the global loss in transcriptional activity is likely higher in WT than *Δvhs* infection due to a slower progression of *Δvhs* infection, relative increases or decreases in transcriptional activity for individual genes remained mostly the same. The only exception was a set of 150 genes that were transcriptionally down-regulated only in WT infection and not *Δvhs* infection (magenta in Fig 3A). These are further analyzed below.

**Figure 3:**
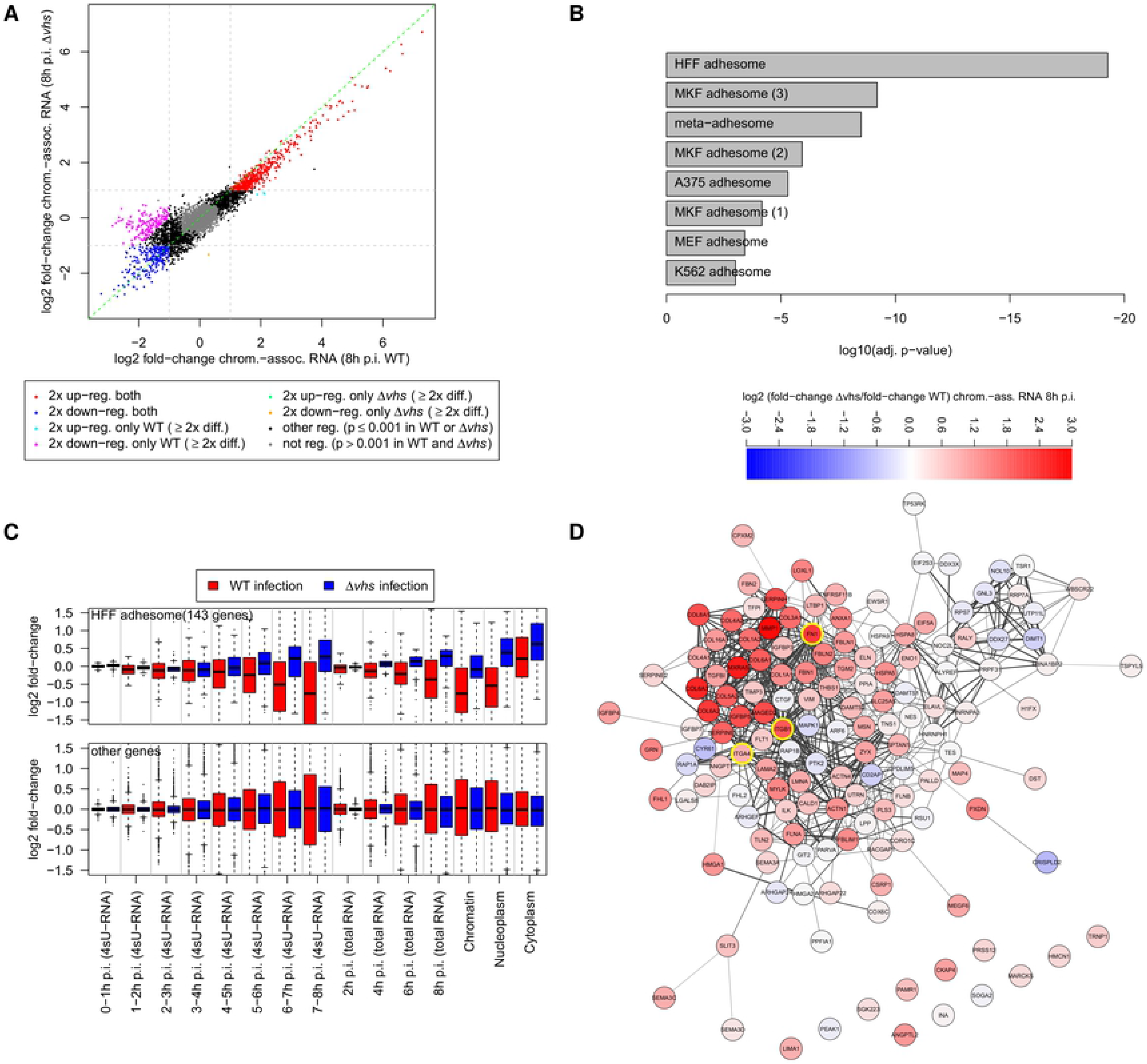
*Vhs*-dependent transcriptional down-regulation of the ECM and integrin adhesome. (A) Scatterplot comparing log2 fold-changes in chromatin-associated RNA at 8h p.i. between WT (x-axis) and *Δvhs* (y-axis) infection. Genes up- (log2 fold-change ≥ 1, adj. p ≤ 0.001) or down-regulated (log2 fold-change ≤ −1, adj. p ≤ 0.001) in both WT and *Δvhs* infection are indicated in red and blue, respectively. Genes transcriptionally down-regulated in a *vhs*-dependent manner (log2 fold-change ≤ −1, adj. p ≤ 0.001 in WT; log2 fold-change > −1 in *Δvhs* infection as well as > 2-fold difference in regulation) are marked in magenta. (B) *vhs*-dependently transcriptionally down-regulated genes are significantly enriched for integrin adhesome components identified in six proteomics studies [29–34] and the meta-adhesome compiled by Horton *et al*. [29]. Barplot shows log10 of multiple testing corrected p-values from Fisher’s exact test. (C) Boxplots showing the distribution of log2 fold-changes in 4sU-RNA, total RNA and subcellular RNA fractions in WT (red) and *Δvhs* infection (blue) for components of the integrin adhesome identified in HFF [32] (top panel) and all other genes (bottom panel). This shows a clear shift in the median of distributions between WT and *Δvhs* infection for the HFF integrin adhesome but not the remaining genes. (D) Protein-protein associations from the STRING database [35] for the HFF integrin adhesome. Colors indicate the log2 ratio between fold-changes in *Δvhs* infection and WT infection (see color bar on top). Red indicates less down-regulation or more up-regulation in *Δvhs* infection than in WT infection and blue the opposite. Yellow borders highlight FN1, the canonical ligand of integrin adhesion complexes, and integrin subunits. The network was visualized with Cytoscape [77].

Notably, 4sU-RNA fold-changes were better correlated to fold-changes in chromatin-associated RNA than to nucleoplasmic or cytoplasmic RNA, while total RNA fold-changes were best correlated to cytoplasmic RNA changes (Fig J in S2 File). This indicates that even with a relatively long 4sU-labeling duration of 60 min, 4sU-RNA to a large degree represents ongoing nascent transcription on the chromatin level. We conclude that fold-changes in chromatin-associated RNA provide an unbiased picture of transcriptional regulation in both WT and *Δvhs* infection.

### *Vhs*-dependent transcriptional down-regulation of the extracellular matrix and integrin adhesome

Differential gene expression analysis on chromatin-associated RNA identified 225 genes (5.4% of all genes) that were significantly down-regulated at the transcriptional level (log2 fold-change ≤ −1, adj. p≤ 0.001) in both WT and in *Δvhs* infection compared to mock (marked blue in Fig 3A). As these genes were characterized by lower poly(A) read-through than non- or up-regulated genes (Fig K in S2 File), their down-regulation cannot be explained by negative effects of read-through transcription on gene expression. Gene Ontology (GO) [25] enrichment analysis for these genes did not yield any statistically significant results, but significant enrichment (adj. p≤0.001) was observed for genes down-regulated by interferon type II according to the INTERFEROME database [26].

Interestingly, a set of 150 genes (3.6% of all genes) was significantly down-regulated (log2 fold-change ≤ −1, adj. p≤ 0.001) in WT but not *Δvhs* infection (marked magenta in Fig 3A, S4 Dataset). Significant v*hs*-dependent down-regulation of these genes was confirmed in nucleoplasmic RNA, 4sU-RNA from 6-7h p.i. onwards and in parts also in total RNA from 6h p.i. onwards (Fig L in S2 File). Similar to *vhs*-independent down-regulated genes, an enrichment for interferon II-down-regulation was observed. Strikingly, however, we also observed a strong functional enrichment for several GO terms (adj. p≤0.001, S5 Dataset), in particular “extracellular matrix (ECM) organization” (>32-fold enriched, adj. *p* < 10^−25^). This included fibronectin (FN1), integrin beta 1 (ITGB1), a subunit of integrin complexes binding fibronectin, and several genes encoding for collagen alpha chains. Enrichment was also observed for “focal adhesion”, i.e. the integrin-containing, multi-protein complexes that anchor the cell to the ECM and connect it to the actin cytoskeleton [27, 28].

Composition of integrin adhesion complexes after induction by their canonical ligand FN1 has been determined by several quantitative proteomics studies in mouse and human cells, including HFF [29–34]. Horton *et al*. consolidated these data into a meta-adhesome of 2,412 proteins found in at least one of six high-quality studies [29]. Adhesome components identified in the individual proteomics studies as well as the meta-adhesome were significantly enriched among genes down-regulated in a *vhs*-dependent manner (Fig 3B, adj. p≤0.001). The highest enrichment was found for the integrin adhesome components identified in HFF (>10-fold enrichment, adj. *p* = 5.3 × 10^−20^). Furthermore, genes of the HFF adhesome (143 genes included in our analysis) showed a systematic shift in regulation between WT and *Δvhs* infection in total RNA, 4sU-RNA and all RNA fractions (Fig 3C). HFF adhesome components tended not to be (or at least less) down-regulated in *Δvhs* infection compared to WT infection, while the remaining genes showed no systematic shift. This shift was already visible from 4-5h onwards in 4sU- and total RNA and when comparing later time points of Δ*vhs* infection to earlier time points of WT infection. Thus, *vhs*-dependent transcriptional down-regulation is not an artefact of comparing different progression stages in the WT and vhs mutant life cycles in 8h p.i. chromatin-associated RNA. When inspecting the protein-protein association network for the HFF adhesome (from the STRING database [35]) the strongest differences between *Δvhs* infection and WT infection were observed in the subnetwork around FN1 and integrin subunits (Fig 3D, Fig M in S2 File).

To investigate whether *vhs*-dependent down-regulation required the *vhs* endonuclease activity, we performed RNA-seq of chromatin-associated RNA at 8h p.i. using a *vhs* single-amino acid mutant (D195N) that no longer exhibited the mRNA decay activity but could still bind to the translation initiation factors eIF4H and eIF4B [36]. For comparison, this was also done for the parental BAC-derived WT virus (WT-BAC) as well as mock, WT and *Δvhs* infection at 8h p.i. (see methods). Interestingly, this confirmed *vhs*-dependent transcriptional regulation of these genes in an independent experiment and demonstrated that it requires *vhs* nuclease activity as fold-changes in D195N infection were extremely well correlated to *Δvhs* infection (Fig N in S2 File). Of note, investigation of RNA-seq read alignments for genomic differences confirmed that the D195N point mutation was the only genome difference of the D195N mutant virus compared to WT-BAC (Fig O in S2 File). This confirms that the D195N mutant expresses the nuclease-null variant of *vhs,* rather than inadvertently no *vhs*. This analysis also confirmed presence of the inactivating insertion in the *Δvhs* mutant [37]. We conclude that components of the integrin adhesome and ECM are transcriptionally downregulated during lytic HSV-1 infection by a *vhs*-nuclease-activity-dependent mechanism.

Gamma-herpesviruses also encode a cytoplasmic mRNA-targeting endonuclease, SOX, which is not homologous to *vhs*. Abernathy *et al*. recently showed that extensive mRNA cleavage by the murine gamma-herpesvirus 68 (MHV68) endoribonuclease muSOX and subsequent Xrn1-mediated mRNA degradation leads to transcriptional repression for numerous genes [17]. In this study, they employed 4sU-seq of WT MHV68 infection and infection with a muSOX-inactivating MHV68 mutant (ΔHS). The same phenomenon was observed for the HSV-1 *vhs* protein when exogenously expressed for 24h. To investigate whether *vhs-*dependent transcriptional down-regulation of the integrin adhesome and ECM components might be mediated by the same mechanism, we compared fold-changes of ΔHS to WT MHV68 infection, on the one hand, and *Δvhs* to WT HSV-1 infection, on the other hand, for both muSOX-dependent genes defined by Abernathy *et al*. and *vhs*-dependent genes defined in this study (Fig P in S2 File). This showed slightly less down-regulation for *vhs-*dependent genes in ΔHS vs. WT MHV68 infection as well as for muSOX-dependent genes in *Δvhs* vs. WT HSV-1 infection than for all other genes in the analysis. However, muSox-dependent genes showed a much more pronounced effect in ΔHS vs. WT MHV68 infection than our *vhs-*dependent genes. Vice versa, *vhs*-dependent genes were much less down-regulated in *Δvhs* vs. WT HSV-1 infection than muSOX-dependent genes. Furthermore, only 14 genes were both muSOX- and *vhs*-dependent. We conclude that, although general mRNA-decay-dependent transcriptional repression may contribute, it is not sufficient to explain the observed differences in transcriptional down-regulation between *Δvhs* and WT HSV-1 infection.

A different explanation for the concerted down-regulation of a set of functionally related genes could be *vhs*-mediated RNA degradation of a key transcriptional regulator. We thus performed motif search in promoters of *vhs*-dependently down-regulated genes but found no significantly enriched novel or known transcription factor binding motifs in proximal promoter regions (−2,000 to +2,000 bp relative to the transcription start site). To recover more distal regulation, we also performed motif search in open chromatin peaks from ATAC-seq data in uninfected cells [20] within 10, 25 or 50kb of *vhs*-dependently down-regulated genes. While this recovered several motif hits for the AP-1 transcription factor, no significant enrichment compared to all identified open chromatin peaks was observed. Interestingly, however, the first *vhs*-dependent gene significantly down-regulated in 4sU-RNA of WT infection at 2-3h p.i. was the ETS transcription factor ELK3, one of three ternary complex factors (TCFs) that act as cofactors of serum response factor (SRF) [38]. SRF has been shown to be vital for focal adhesion assembly in embryonic stem cells [39]. TCF-dependent genes identified from simultaneous knockouts of all three TCFs as well as SRF targets from ChIP-seq have previously been determined in mouse embryonic fibroblasts (MEFs) [40]. Though we found no significant enrichment for TCF-dependent genes or TCF-dependent SRF targets, SRF targets in general were significantly enriched (~2.25-fold) among *vhs*-dependent genes (*p* = 3.9 × 10^−5^). Nevertheless, only 42 (28%) of *vhs*-dependent genes were SRF targets and 93% of SRF targets were not *vhs*-dependent in our study, thus other regulatory mechanisms have to be involved.

Pheasant *et al*. observed large differences regarding the extent of *vhs*-induced loss in total RNA levels between different cellular genes at 12h WT infection [18]. Using qRT-PCR, they showed that this reduction was *vhs*-dependent based on two sets of genes that exhibited either high (COL6A, MMP3, MMP1) or low reduction (GAPDH, ACTB, RPLP0) in total RNA levels in WT infection. As Actinomycin D treatment confirmed similar stability of corresponding mRNAs, they concluded that these differences were not due to differences in transcription rates or mRNA stability between these genes but rather due to differences in the susceptibility of the respective transcripts to *vhs* cleavage activity. We noted that two (COL6A1, MMP1) of the PCR-confirmed highly *vhs*-sensitive genes belonged to our set of genes transcriptionally downregulated in a *vhs*-dependent manner. The third gene (MMP3), though originally not included in our analysis due to its proximity to nearby genes, is also involved in ECM organization. Moreover, genes defined as efficiently depleted during WT infection by Pheasant *et al*. (log2 fold-change in total RNA at 12h p.i. WT infection < −5) were significantly enriched for ECM organization (>3-fold, adj. *p* = 7.4 × 10^−7^). We thus hypothesized that a significant fraction of highly *vhs*-sensitive genes identified by Pheasant *et al*. may actually be transcriptionally down-regulated in a *vhs*-dependent manner. To test this hypothesis, we performed differential gene expression analysis in chromatin-associated RNA for all human genes and identified an extended set of *vhs*-dependently transcriptionally down-regulated genes (578 genes, which now also included MMP3, S6 Dataset). The additional *vhs*-dependent genes were also significantly enriched for focal adhesion and ECM organization (>11-fold enriched, adj. *p* < 10^−23^). Both original and additional *vhs*-dependent genes were strongly enriched among efficiently depleted genes determined by Pheasant *et al*. (4.2 - 6.8-fold enrichment, *p* < 10^−27^) and were among the most significantly down-regulated genes in total RNA at 12h p.i. in WT infection (Fig Q in S2 File). We conclude that *vhs*-dependent transcriptional down-regulation notably contributes to reduced total mRNA levels of the respective genes later on in WT HSV-1 infection and thereby explains the previously observed differences in “*vhs* activity” on the mRNA levels of these genes.

### A common core of up-regulated genes in WT and *Δvhs* infection

Analysis of chromatin-associated RNA identified a set of 462 genes that were significantly up-regulated in both WT and *Δvhs* infection (log2 fold-change ≥ 1, adj. *p* ≤ 0.001, marked red in Fig 3A). Only 3 genes were up-regulated in WT but not or 2-fold less in *Δvhs* infection. Thus, transcriptional up-regulation during HSV-1 infection is independent of *vhs*. Clustering analysis of *vhs*-independent up-regulated genes identified four subgroups that were distinguished mostly by how strongly and early in infection they were up-regulated (Fig 4A, S7 Dataset). In particular, a set of 24 genes (marked orange in Fig 4A) was up-regulated both very early and strongly in WT and *Δvhs* infection, with up-regulation of 21 of these genes (91.7%) detectable in total RNA at 6h p.i. or earlier in both WT and *Δvhs* infection (Fig R in S2 File). Not surprisingly, several of these genes (e.g. RASD1, NEFM, NPTX2) had previously been identified as highly up-regulated in HSV-1 infection by microarray analysis on total RNA [41, 42] and 10 were significantly up-regulated in total RNA at 12h p.i. WT infection in the Pheasant *et al*. data [18]. Up-regulation of all orange and blue cluster genes was also confirmed in 4sU-RNA (Fig S in S2 File). No enrichment for GO terms was observed either for individual clusters or all up-regulated genes, however the green and orange cluster were enriched for interferon type I-up-regulated genes (adj. *p* = 1.68 × 10^−5^ and adj. p=0.0019 for the green and orange cluster, respectively). Notably, 50% of genes in the orange cluster were up-regulated by type I interferons (>4.5-fold enrichment).

**Figure 4:**
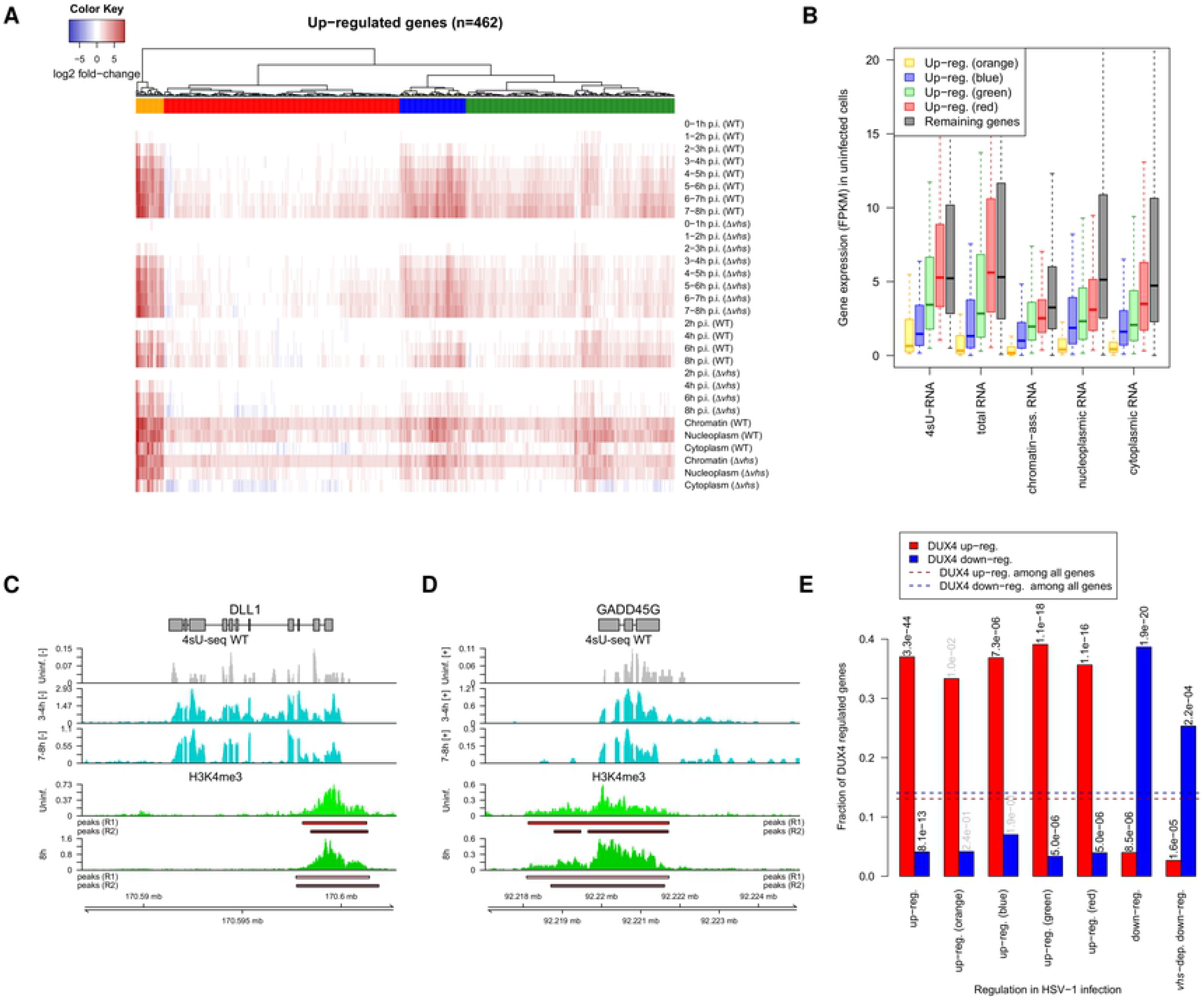
*vhs*-independent transcriptional up-regulation of lowly expressed genes. (A) Heatmap of log2 fold-changes in 4sU-RNA, total RNA and subcellular RNA fractions for genes up-regulated in both WT and *Δvhs* infection (marked red in Fig 3A). Genes were clustered according to Euclidean distances and Ward’s clustering criterion (see methods). Four clusters were obtained at a distance threshold of 30 and are indicated by colored rectangles (orange, blue, green, red). (B) Boxplots of the distribution of expression values (FPKM) in uninfected cells from 4sU-RNA, total RNA and subcellular RNA fractions show low or no expression of strongly up-regulated genes (orange cluster) in uninfected cells compared to other up-regulated clusters (blue, green, red) and remaining genes. (C-D) Strongly up-regulated genes with low expression in uninfected cells, such as DLL1 (C, negative strand) and GADD45G (D, positive strand), are already primed for up-regulation by H3K4me3 marks at their promoters. Tracks show read coverage (normalized to total number of mapped human reads; averaged between replicates) in uninfected and WT 4sU-RNA for selected time points (gray and cyan, top 3 tracks) and H3K4me3 ChIPmentation in uninfected cells and at 8h p.i. WT infection (green, bottom 2 tracks). Peaks identified in each replicate are shown separately below H3K4me3 read coverage tracks. Gene annotation is indicated on top. Boxes represent exons and lines introns. Genomic coordinates are shown on the bottom. For 4sU-seq data only read coverage on the same strand as the gene is shown (+ = positive strand, − = negative strand). H3K4me3 ChIPmentation is not strand-specific. (E) Barplots showing the fraction of transcriptionally regulated genes in HSV-1 infection that are either up- (red) or down- (blue) regulated by doxycycline-inducible DUX4 [49]. Results are shown separately for genes up-regulated in both WT and *Δvhs* infection, the four clusters of up-regulated genes, genes down-regulated in both WT and *Δvhs* infection as well as genes down-regulated in a *vhs*-dependent manner in WT infection. Horizontal dashed lines indicate the fraction of all analyzed genes regulated by DUX4. Numbers on top of bars indicate p-values (corrected for multiple testing) for a Fisher’s exact test comparing the fraction of DUX4 up- or down-regulated genes between each group of HSV-1 regulated genes to the background of all genes (black: adj. p≤0.001, gray: not significant).

One characteristic feature of up-regulated genes in general and the orange cluster in particular was their low level of gene expression in uninfected cells (Fig 4B). Notably, 71% of genes in the orange cluster were not or only very lowly expressed (total RNA FPKM ≤1) in uninfected cells compared to 8% of all genes (Fisher’s exact test *p* < 10^−13^). In total, 76 (17, 22, 32, 5 from the orange, blue, green and red cluster, respectively) up-regulated genes (16.5%) were poorly expressed in uninfected cells. HSV-1 induced up-regulation of genes not normally expressed has previously been reported for human alpha globin genes (HBA1, HBA2), which are normally only expressed in erythroid cells [43]. RNA-seq analysis of these two duplicated genes is complicated by their high sequence similarity (>99% on coding sequence, 5’ UTRs and upstream of promoter [44]), as most reads can be mapped equally well to both genes and their promoter regions. Nevertheless, our data clearly confirmed that at least one of the two alpha globin genes is transcribed during HSV-1 infection as early as 2-3h p.i. and translated into protein at least from 4h p.i. (according to our previously published Ribo-seq data [19]). Our analysis suggests that similar up-regulation from no or low expression is observed for a number of other cellular genes. Since we used relatively strict criteria to exclude genes that only appeared to be expressed during infection due to read-in transcription, we also investigated more lenient criteria to identify the extent of induction for genes that are not expressed prior to infection (see methods for details). These criteria applied to 17 of the up-regulated genes (e.g. DLL1) and an additional 33 genes not included in our previous analysis. Manual inspection of these 33 genes confirmed clear transcriptional up-regulation only for 13 genes (IRF4, RRAD, FOSB, ARC, CA2, DIO3, DLX3, GBX2, ICOSLG, MAFA, MAFB, NGFR, PCDH19). Of these, 6 and 8 were up-regulated by type I and II interferons, respectively. In summary, only a small fraction of genes not expressed in uninfected fibroblasts is induced by HSV-1 infection.

To start investigating how rapid up-regulation of these genes might be achieved, we performed ChIPmentation [45] of H3K4me3 histone marks (2 replicates each in uninfected cells and at 8h p.i. WT infection). H3K4me3 has been reported to regulate assembly of the preinitiation complex for rapid gene activation [46]. Furthermore, a bivalent chromatin modification pattern combining H3K4me3 and H3K27me3 has been described in embryonic stem (ES) cells, which serves to keep silenced developmental genes poised for activation [47]. Across all 4 samples, we identified 32,601 unique non-overlapping peak regions, which were strongly enriched around gene promoters (Fig T in S2 File, S8 Dataset). In total, 98.7% of analyzed genes exhibited H3K4me3 peaks around the promoter in both replicates of uninfected cells. Notably, this also applied to 21 of the 24 genes in the orange cluster (87.5%, see Fig 4C,D for examples). Only NPTX1 and NPTX2 showed no significant H3K4me3 promoter peak in either replicate of uninfected cells. Both showed peaks in at least one replicate of infected cells (Fig U in S2 File). In total, 97.8% of all up-regulated genes and 92.1% of up-regulated genes that were not or lowly expressed in uninfected cells (total RNA FPKM ≤1) showed significant peaks in both replicates of uninfected cells. In summary, this indicates strong, early, *vhs-*independent transcriptional up-regulation of a small number of poorly expressed genes which are already poised for expression by H3K4me3 marks at their promoters.

Recently, Full *et al*. reported that germline transcription factor double homeobox 4 (DUX4) and several of its targets were highly upregulated by HSV-1 infection [48]. We thus compared genes up- or down-regulated by doxycycline-inducible DUX4 [49] with genes transcriptionally regulated in HSV-1 infection (Fig 4E). We found that HSV-1 up-regulated genes were significantly (Fisher’s exact test, adj. p≤0.001) enriched for DUX4 up-regulation and HSV-1 down-regulated genes were significantly enriched for DUX4 down-regulation. Notably, the fraction of genes up-regulated by DUX4 was similar (~36%) for all clusters of HSV-1 up-regulated genes, independent of their expression in uninfected cells. Interestingly, however, genes transcriptionally down-regulated in HSV-1 infection in a *vhs*-mediated manner were less enriched for DUX4-mediated down-regulation than genes for which transcriptional down-regulation was independent of *vhs.* Moreover, enrichment for adhesome components was more pronounced among *vhs*-dependent genes not down-regulated by DUX4 than among those down-regulated by DUX4. Thus, while DUX4 is a major transcriptional regulator in HSV-1 infection, it is not responsible for *vhs*-mediated down-regulation of the integrin adhesome.

### *Vhs*-dependent transcriptional down-regulation impacts on cellular protein levels

To investigate how changes in total RNA levels and transcription alter protein levels in infected cells, we performed a Tandem Mass Tag (TMT)-based quantitative proteomic analysis of WT- and Δ*vhs*-infected HFF at 0 and 8h p.i. (n=3 replicates). In total, 7,943 proteins were identified (S9 Dataset). No filtering based on read-in transcription was performed, as read-through transcripts are neither exported nor translated [19, 20]. Protein fold-changes were poorly correlated to fold-changes in total, 4sU- or subcellular RNA fractions (*r*_*s*_ ≤ 0.21, Fig V in S2 File) and generally tended to be less pronounced. Both observations are consistent with the higher stability of proteins compared to mRNAs (~5 times more stable in mouse fibroblasts [50]), thus changes in *de novo* transcription and total RNA levels commonly take >8h to significantly impact on protein levels. Consequently, protein fold-changes were very well correlated between WT and Δ*vhs* infection (Fig 5A, *r*_*s*_ = 0.96) and only few cellular proteins showed a significant difference between WT and Δ*vhs* infection. Due to the less pronounced changes, we determined differentially expressed proteins with a >1.5-fold change (adj. *p* ≤ 0.001, Fig 5A). Most differentially expressed proteins were concordantly regulated either down (1,444 genes, 73%) or up (499 genes, 25.3%) in both WT and Δ*vhs* infection. It should be noted that, similar to RNA-seq data, protein fold-changes only represent relative changes in the presence of a global loss in cellular protein levels. Thus, some up-regulated proteins may simply be less/not down-regulated compared to most other proteins.

**Figure 5:**
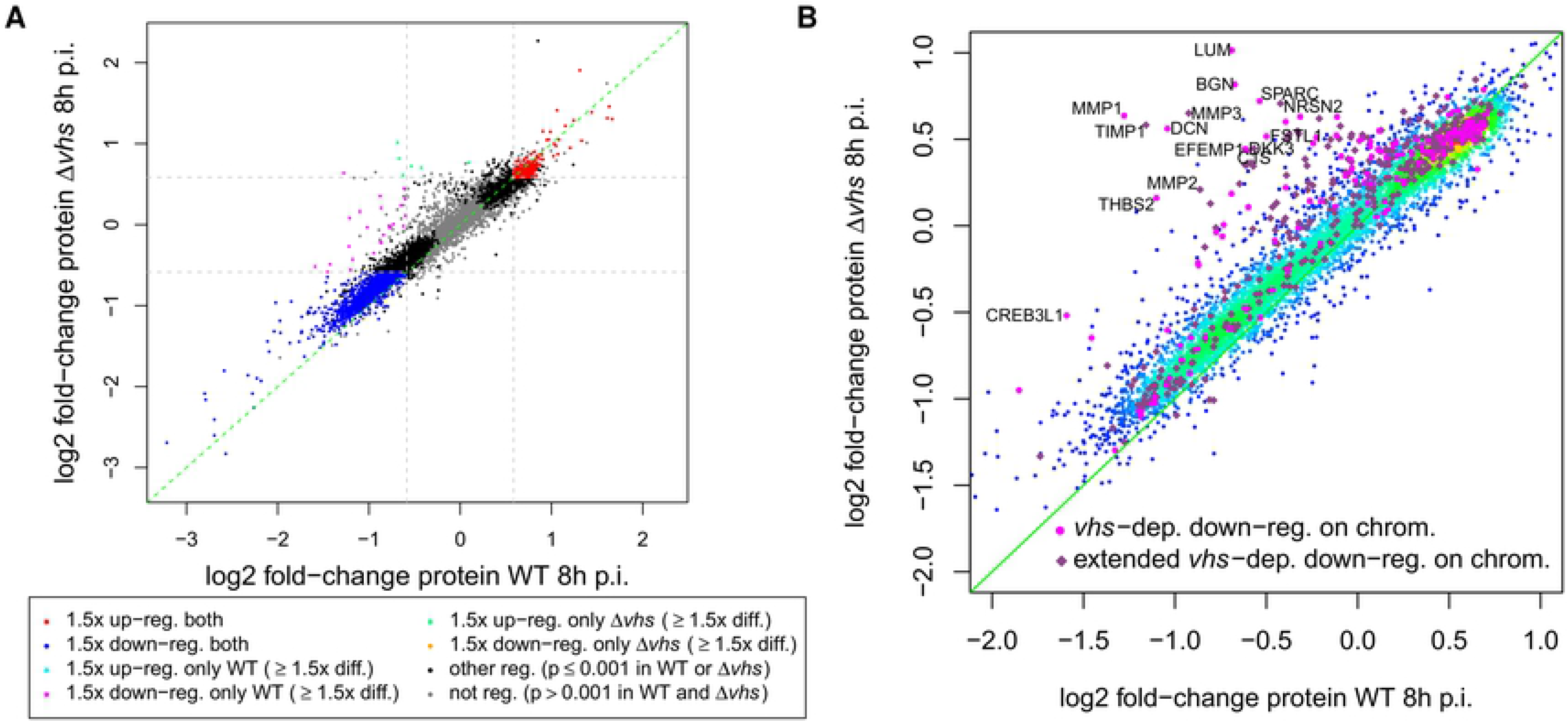
Impact of HSV-1 infection on protein levels. (A) Scatterplot comparing log2 fold-changes in protein levels at 8h p.i. between WT (x-axis) and *Δvhs* (y-axis) infection. Up- or down-regulated proteins (≥ 1.5-fold change, adj. p ≤ 0.001) in both WT and *Δvhs* infection are indicated in red and blue, respectively. Proteins down-regulated in a *vhs*-dependent manner (≥ 1.5-fold down-regulated, adj. p ≤ 0.001 in WT; <1.5-fold down-regulated in *Δvhs* infection as well as >1.5-fold difference in regulation) are marked in magenta. Green indicates proteins that are up-regulated in *Δvhs* infection but not in WT infection with a >1.5-fold difference in fold-changes. (B) Scatterplot comparing log2 fold-changes in protein levels at 8h p.i. between WT (x-axis) and *Δvhs* (y-axis) infection. Points are color-coded according to density of points: from red = high density to blue = low density. Pink and violet points represent genes that are down-regulated in chromatin-associated RNA in a v*hs*-dependent manner. Here, pink indicates genes included in our original analysis and violet genes that were additionally identified from the differential gene expression analysis on all genes. Gene symbols are shown for genes with a ≥ 2-fold increase in protein fold-changes in *Δvhs* infection compared to WT infection.

Concordantly, down-regulated proteins were significantly (adj. p≤0.001, S10 Dataset) enriched for a number of GO terms, including “nucleotide-sugar biosynthetic process” (>77-fold enriched), “canonical glycolysis” (>15-fold), “viral budding” (>9-fold) and “activation of MAPK activity” (>4-fold). Interestingly, meta-adhesome (but not HFF adhesome) components were also significantly enriched (>1.9-fold), indicating that concordantly down-regulated proteins interact with the core adhesome, rather than are a part of it. Interestingly, concordantly up-regulated proteins were highly enriched for mitochondrial proteins (>7-fold, 186 proteins), but significantly depleted of meta-adhesome components. Of the 19 genes significantly up-regulated in total RNA in both WT and Δvhs infection, 5 were also up-regulated at protein level. This included 4 genes that were up-regulated in chromatin-associated RNA (RASD1, SNAI1, CBX4, ITPR1).

Thus, transcriptional up-regulation does have a small but significant effect on protein levels by 8h p.i. Only few genes showed a significant differential effect (24 down-regulated in WT only, 6 up-regulated in Δ*vhs* only). Strikingly, the 24 proteins down-regulated in a *vhs*-dependent manner were strongly enriched for HFF adhesome components (>12-fold) and ECM organization (>23-fold). Accordingly, 9 of 14 (64%) proteins down-regulated only in WT infection and included in our RNA-seq analysis were down-regulated in chromatin-associated RNA in a *vhs*-dependent manner. An analysis of protein fold-changes for all *vhs*-dependently transcriptionally down-regulated genes (including our extended set) showed that a significant number of respective proteins tended to be either less down-regulated or (relatively) more up-regulated in Δ*vhs* infection than in WT infection (Fig 5B). Many of these were components of the integrin adhesome or were involved in ECM organization. Thus, *vhs*-dependent transcriptional down-regulation impacts protein levels of the respective genes already by 8h p.i.

## Discussion

HSV-1 infection drastically alters host RNA metabolism at all levels by impairing host mRNA synthesis, processing, export and stability. Here, we differentiate and quantify their individual contributions to the RNA expression profile by combining RNA-seq of total, newly transcribed (4sU-)RNA and subcellular RNA fractions in WT and Δ*vhs* infection. We developed a mathematical model to quantify both the loss of transcriptional activity and the changes in *vhs* nuclease activity based on the correlations between RNA half-lives and total RNA fold-changes during the first 8h of infection. This depicted a drop in transcriptional activity down to 10-20% of the original level by 8h p.i., consistent with the well-described general loss of Pol II from host chromatin [14, 15]. Comparison of WT and Δ*vhs* infection confirmed a rapid increase in *vhs*-dependent degradation, with 20-30% of all cellular mRNAs degraded per hour by 2h p.i., consistent with the well-described role of *vhs* upon viral entry. While *vhs* activity did not further rise from 4h p.i., it was constantly maintained until 8h p.i. The kinetics of the viral life cycle are incorporated in our mathematical model via the functions describing *vhs* activity and cellular transcriptional activity. As *vhs* activity and cellular transcriptional activity cannot be estimated simultaneously in WT infection, we estimated the development of host transcriptional activity in WT from *Δvhs* infection. Considering the slower progression of *Δvhs* infection, we may have thus underestimated the drop in transcriptional activity in WT infection. However, if transcriptional activity drops even faster and further in WT infection, *vhs* activity would have to increase even faster and to higher levels to explain the observed negative correlations between RNA half-lives and total RNA fold-changes in WT infection. It is important to note that our findings do not contradict previous reports on the post-transcriptional down-regulation of *vhs* activity by its interaction with VP16 and VP22 [9–11]. As previously already noted [9], counter-regulation of *vhs* activity is not complete, but VP16 and VP22 clearly serve to prevent a further detrimental increase in *vhs* activity during infection. Their activity thus explains the plateau we observed for *vhs* activity despite substantially increasing *vhs* protein levels. Moreover, application of our model to total RNA fold-changes at 12h p.i. WT infection from the study of Pheasant *et al*. confirmed rapid deactivation of *vhs* between 8 and 12h p.i.

Pheasant *et al*. also noted that *vhs*-dependent reduction in total RNA levels varied widely between genes at 12h p.i. and hypothesized that this might indicate differences in susceptibility to *vhs*-mediated degradation between transcripts [18]. Furthermore, they excluded an influence of basal transcription rates and RNA half-lives for the three genes whose high *vhs*-sensitivity they confirmed by PCR. However, we here show that all three genes they chose are actually transcriptionally down-regulated in a *vhs*-dependent manner. Together with the effects of *vhs* on RNA stability, this translates into a significant reduction in the corresponding protein levels by 8h p.i. Accordingly, results from these three genes cannot be extrapolated to genes down-regulated in total RNA *only* through *vhs*-mediated RNA decay. Instead, our mathematical model suggests that gene-specific differences in mRNA half-lives substantially shape the variability in total mRNA changes between genes at least until 8h p.i. This does not exclude a contribution of other factors, e.g. *vhs*-induced nuclear retention of cellular mRNAs shown by Pheasant *et al*. [18] or differences in translation rates between different mRNAs (and thus translation-initiation-dependent mRNA cleavage by the *vhs* protein), which we did not consider in our model. In particular, *vhs*-dependent transcriptional down-regulation contributes substantially to the reduction in total RNA levels for the respective genes (Fig Q in S2 File, Fig W in S2 File). Furthermore, a recent study identified a set of 74 genes that escape degradation by four herpesviral endonucleases, including *vhs* [51]. Almost all of these genes were excluded from our analysis due to low expression (87%), read-in transcription (7%) or proximity to nearby genes (3%). Two genes, however, which were not excluded, (C19orf66, ARMC10) indeed did not show any significant change in any of our data. Thus, while we cannot exclude that some transcripts are less susceptible to *vhs*-mediated decay than others, we can conclude that strong reductions in total mRNA levels are not necessarily a consequence of increased susceptibility of individual transcripts to *vhs*-mediated RNA cleavage.

In contrast to total cellular RNA changes, fold-changes in newly transcribed and, in particular, chromatin-associated RNA were highly similar between WT and Δ*vhs* infection. Although we performed the total RNA- and 4sU-seq time-courses for both viruses in separate experiments, the high correlation of 4sU-RNA fold-changes confirmed that the obtained RNA-seq data could indeed be directly compared. Furthermore, it enabled us to decipher gene-specific transcriptional regulation that is either dependent or independent of *vhs*. While the analysis of chromatin-associated RNA eliminated the bias originating from *vhs* activity and the global loss in transcription, read-in transcription leading to seeming, but non-functional, induction of genes has to be taken into account in all gene expression profiling studies independent of the type of profiled RNA. By excluding genes with evidence of read-in transcription from our analysis, we ascertained that all identified induced genes represent true up-regulation and not artefacts from read-in transcription. Notably, while most strongly up-regulated genes identified in our study have been reported in previous studies on HSV-1-induced differential host expression (e.g. RASD1 [18, 41, 52, 53]), several previously reported genes, which were thought to be induced by HSV-1, are actually only seemingly induced due to read-in transcription, e.g. ZSCAN4 [41, 54], SHH [52] and FAM71A [18].

Around 30% of all up-regulated genes and 50% of the most strongly up-regulated genes (orange cluster) were up-regulated by type I interferons (IFN). Moreover, DUX4 was confirmed as a major transcriptional regulator in both WT and Δ*vhs* infection for both up- and down-regulated genes (37% of up-regulated genes were previously found to be up-regulated by DUX4 and 39% of down-regulated genes were down-regulated by DUX4, Fig 4E). Although there was some overlap between DUX4 and IFN-induced genes amongst the HSV-1-induced genes, it was not significantly higher than expected at random. Interestingly, the DUX4 up-regulated gene TRIM43 was recently identified as a herpesvirus-specific antiviral factor independent of the type I interferon response [48], suggesting that DUX4-mediated regulation in HSV-1 infection may represent an alternative pathway which augments the host intrinsic immune response.

A key finding of our study is the *vhs*-dependent, transcriptional down-regulation of proteins involved in the integrin adhesome and ECM organization, which required *vhs* nuclease activity. Suppression of ECM protein synthesis during HSV-1 infection has already been shown over 30 years ago for the canonical integrin ligand FN1, type IV procollagen and thrombospondin [55]. Recently, this was confirmed for a few other ECM components in human nucleus pulposus cells in both lytic and latent HSV-1 infection [56]. *Vhs*-dependency of down-regulation was previously reported for FN1 [57], but was ascribed to the effect of *vhs* on FN1 RNA stability. This further highlights the fallacy in ascribing all *vhs*-dependent effects on total RNA levels directly to *vhs*-mediated RNA decay. In contrast, our data demonstrates that *vhs*-dependent down-regulation of many genes is augmented by *vhs*-dependent repression of transcription. Notably, while *vhs*-dependent down-regulation of the ECM and adhesome can largely be confirmed in total RNA, it is challenging to distinguish from *vhs*-mediated mRNA degradation (Fig W in S2 File). The transcriptional effects only become obvious when analyzing chromatin-associated RNA.

Interestingly, transcriptional down-regulation of ECM and integrin adhesome genes was dependent on the nuclease activity of *vhs*. Recently, muSOX-mediated RNA decay was reported to trigger wide-spread transcriptional repression at late times of lytic MHV68 infection [17]. While HSV-1 *vhs* activity also triggered this phenomenon within 24h of expression, the cellular genes transcriptionally regulated in a *vhs*-dependent manner during the first 8h of HSV-1 infection showed little overlap to the genes affected by the transcriptional effects of muSOX-induced RNA degradation.

An alternative explanation for the *vhs*-dependent repression of such a functionally connected cellular network of genes is that *vhs* nuclease activity results in a rapid depletion of transcripts of key, short-lived cellular transcription factor(s) governing these genes. It is unclear, however, why only a single or very small number of transcription factors would suffer so dramatically from *vhs* nuclease effects. It is indeed surprising that *vhs*-mediated mRNA degradation does not cause a similarly pronounced dysregulation downstream of short-lived transcription factors involved in other processes. However, the surprisingly high correlation between fold-changes in WT and Δ*vhs* infection observed in chromatin-associated RNA excludes gross global effects of mRNA degradation of cellular transcriptional factors. Furthermore, no enriched known or novel transcription factor binding motif could be identified in both proximal promoter regions or more distal open chromatin regions identified by ATAC-seq. Promoter analysis applied to all expressed HFF adhesome genes also identified only one significant motif in less than 6% of genes, suggesting that there is no single key transcriptional regulator for the integrin adhesome. Nevertheless, *vhs* may still directly interact with or target a major cellular transcription factor that governs the expression of the integrin adhesome and ECM via distal enhancers. Notably, ELK3, a TCF co-factor of SRF, was down-regulated in a *vhs*-dependent manner early on in infection. While TCF-dependent genes were not enriched among *vhs*-dependent genes, a ~2-fold enrichment of SRF targets was observed. Since TCF-dependent genes were determined by triple knockouts of all three TCFs, not all TCF-dependent genes likely depend on ELK3. Thus, ELK3-dependent reduced recruitment of SRF could still play a role. In addition, post-transcriptional processes have been linked to transcriptional control of focal adhesions and may thus also be relevant for *vhs*-dependent down-regulation. For instance, Rho signaling can result in nuclear translocation of the SRF co-factor MRTF-A and prevention of this translocation results in lower expression of cytoskeletal/focal adhesion proteins [58]. Furthermore, in keratinocytes nuclear actins lead to down-regulation of a number of adhesion proteins [59], such as ITGB1 and MYL9, which were also down-regulated in a *vhs*-dependent manner in HSV-1 infection.

Untangling the molecular mechanisms underlying specific *vhs*-mediated down-regulation of the integrin adhesome and ECM will be difficult without knowledge of the responsible cellular transcription factor(s) and confounded by the pleotropic effects of *vhs* nuclease activity. Nevertheless, we could show that *vhs*-dependent transcriptional down-regulation has a clear impact on protein levels already by 8h p.i., as confirmed by quantitative whole cell proteomics. Proteins with strong *vhs*-dependent reduction at 8h p.i. include matrix metallopeptidases MMP1-3, which are involved in degradation of ECM proteins, their inhibitor TIMP1 as well as other MMP-up-regulating or -interacting proteins (LUM, SPARC, THBS2).

In summary, our analyses provide a quantitative picture of the molecular mechanisms that govern profound alterations in the host cell transcriptome and proteome during lytic HSV-1 infection.

## Materials and Methods

### Cell culture and infections

Human fetal foreskin fibroblasts (HFF) were purchased from ECACC (#86031405) and cultured in DMEM with 10% FBS Mycoplex and 1% penicillin/streptomycin. HFF were utilized from passage 11 to 17 for all high-throughput experiments. This study was performed using WT HSV-1 strain 17 (data taken from previous studies [19, 20]) and its *vhs*-inactivated mutant (*Δvhs*) [60]. Virus stocks were produced in baby hamster kidney (BHK) cells (obtained from ATCC) as described [19]. HFF were infected with HSV-1 24h after the last split for 15 min (for total RNA-seq, 4sU-seq and RNA-seq of subcellular fractions) or 1h (for RNA-seq of chromatin-associated RNA including the *vhs* D195N mutant), at 37°C using a multiplicity of infection (MOI) of 10. Subsequently, the inoculum was removed and fresh media was applied to the cells.

The *vhs* D195N mutant virus was constructed via *en passant* mutagenesis [61]. Mutagenesis templates were generated using PCR primers GTATATCTGGCCCGTACATCGATCT and GGTCAGTGTCCGTGGTGTACACGTACGCGACCGTGTTGGTGTGATAGAGGTTG GCGCAGGCATTGTCCGCCTCCAGCTGACCCGAGTTAAAGGATGACGACGATAA GTAGGG to amplify the kanamycin resistance cassette flanked by Isce-I restriction sites from vector pEP-Kan. Additional homologies for recombination were added to this product by PCR using primers GGTCAGTGTCCGTGGTGTAC and TTCTGTATTCGCGTTCTCCGGGCCCTGGGGTACGCCTACATTAACTCGGGTCAGCTGGAGGCGGACAATGCCTGCGCCAACCTCTATCACGTATATCTGGCCCGTACATCGATCT before electroporation into *Escherichia coli* strain GS1783 containing the pHSV(17+)Lox BAC [62]. BAC DNA was purified using the NucleoBond BAC 100 kit (Macherey-Nagel #740579) and transfected for virus reconstitution into BHK-21 cells with Lipofectamine 3000 (ThermoFisher #L3000-075).

### Preparation of RNA

Sample preparation for 4sU-seq in *Δvhs* infection was performed as reported previously for WT HSV-1 [19]. In brief, 4-thiouridine (4sU) was added to the cell culture medium for 60 min at −1, 0, 1, 2, 3, 4, 5, 6, or 7h p.i. (2 × 15-cm dishes per condition) during *Δvhs* infection to a final concentration of 500 μM (n=2 replicates). Subsequently, the medium was aspirated and the cells were lysed with Trizol (Invitrogen). Total RNA and newly transcribed RNA fractions were isolated from the cells as described previously [24]. In an independent experiment, subcellular RNA fractions (cytoplasmic, nucleoplasmic and chromatin-associated RNA) in mock and 8h p.i. of WT and *Δvhs* infection were prepared as previously described (n=2 replicates) [20]. To assess the role of *vhs* nuclease activity in regulation of ECM and integrin adhesome genes, chromatin-associated RNA in mock, WT, *Δvhs, vhs* D195N and WT-BAC infection at 8h p.i. (n=2 replicates) was prepared.

### Library preparation and RNA sequencing

Sequencing libraries were prepared using the TruSeq Stranded Total RNA kit (Illumina). rRNA depletion was performed after DNase treatment for total RNA and all subcellular RNA fractions using Ribo-zero but not 4sU-RNA samples. Sequencing of 75bp paired-end reads was performed on a NextSeq 500 (Illumina) at the Core Unit Systemmedizin (Würzburg).

### H3K4me3 ChIPmentation

The full description of H3K4me3 ChIPmentation is included in S11 Text.

### Preparation of samples for proteomic analysis

HFF were infected with WT HSV-1 or its *vhs*-inactivated mutant for 8h at an MOI of 10. Infections were conducted in triplicate, with 4 uninfected controls (10 samples in total). Washed cells were snap-frozen in liquid nitrogen. Cells were lysed in by resuspending in 100μL 2% SDS/50mM TEAB pH 8.5 followed by 10mins (30s on/off duty cycle) sonication in a bioruptor sonicator (Diagenode). Lysates were quantified by BCA assay and 50μg of each sample was reduced and alkylated with 10mM TCEP and 40mM Iodoacetamide for 20 minutes at room temperature in the dark. Samples were made up to 500uL with 8M urea 50mM TEAB and applied to 30kDa Vivacon centrifugal ultrafiltration devices (Sartorius). Samples were concentrated according to the manufacturer’s instructions. Samples were resuspended and concentrated in 8M urea buffer a further 3 times to remove residual SDS. There were a further 3 washes with digestion buffer (0.5% Sodium deoxycholate 50mM TEAB) before samples were resuspended in approximately 50uL digestion buffer with 1ug Trypsin (Proteomics grade, Thermo Fisher). Filter units were then incubated in at 37 degrees overnight in a box partially filled with water to reduce evaporation. Peptides were recovered into a fresh tube by centrifugation and a further wash with 50uL digestion buffer. SDC was removed from each sample by precipitation with the addition of formic acid and two-phase partitioning with ethyl acetate. Peptides were then dried under vacuum. For TMT labelling samples were resuspended in 42uL 100mM TEAB and 0.4mg of each TMT reagent in 18uL anhydrous acetonitrile was added, vortexed to mix and incubated at room temperature for 1 hour. A small aliquot of each sample was analyzed by LC-MS to confirm labelling efficiency and samples were pooled 1:1 according to the total TMT reporter intensity in these QC runs. The pooled sample was then acidified and subjected to SPE clean-up using 50mg tC18 cartridges (Waters) before drying under Vacuum.

### Basic pH Revered Phase fractionation

Samples were resuspended in 40μL 200mM Ammonium formate pH10 and transferred to a glass HPLC vial. BpH-RP fractionation was conducted on an Ultimate 3000 UHPLC system (Thermo Scientific) equipped with a 2.1 mm × 15 cm, 1.7μ Kinetex EVO column (Phenomenex). Solvent A was 3% ACN, Solvent B was 100% ACN, solvent C was 200 mM ammonium formate (pH 10). Throughout the analysis solvent C was kept at a constant 10%. The flow rate was 400 μL/min and UV was monitored at 280 nm. Samples were loaded in 90% A for 10 min before a gradient elution of 0– 10% B over 10 min (curve 3), 10-34% B over 21 min (curve 5), 34-50% B over 5 mins (curve 5) followed by a 10 min wash with 90% B. 15s (100μL) fractions were collected throughout the run. Fractions containing peptide (as determined by A280) were recombined across the gradient to preserve orthogonality with on-line low pH RP separation. For example, fractions 1, 25, 49, 73, 97 are combined and dried in a vacuum centrifuge and stored at −20°C until LC-MS analysis.

### Mass Spectrometry

Samples were analysed on an Orbitrap Fusion instrument on-line with an Ultimate 3000 RSLC nano UHPLC system (Thermo Fisher). Samples were resuspended in 10μL 5% DMSO/1% TFA. 5μL of each fraction was Injected. Trapping solvent was 0.1% TFA, analytical solvent A was 0.1% FA, solvent B was ACN with 0.1% FA. Samples were loaded onto a trapping column (300μm x 5mm PepMap cartridge trap (Thermo Fisher)) at 10μL/min for 5 minutes. Samples were then separated on a 50cm × 75μm i.d. 2μm particle size PepMap C18 column (Thermo Fisher). The gradient was 3-10% B over 10mins, 10-35% B over 155 minutes, 35-45% B over 9 minutes followed by a wash at 95% B for 5minutes and requilibration at 3% B. Eluted peptides were introduced by electrospray to the MS by applying 2.1kV to a stainless steel emitter (5cm × 30μm (Thermo Fisher)). During the gradient elution, MS1 spectra were acquired in the orbitrap, CID-MS2 acquired in the ion trap. SPS isolated MS2 fragment ions were further fragmented using HCD to liberate reporter ions which were acquired in the orbitrap (MS3).

### Data Processing

Raw files were searched using Mascot (Matrix Science) from within Proteome Discoverer Ver 2.1 (Thermo Fisher) against the uniport human database with appended common contaminants and uniport HSV reference proteome. PSM FDR was controlled at 1% using Mascot Percolator. The reporter ion intensities of proteins with a High (1%) and Medium (5%) FDR were taken and subjected to LIMMA t-test in R. P-values were adjusted for multiple testing using the method by Benjamini and Hochberg [63]. Proteins with extremely high standard deviation between replicates in (>99 percentile) in either WT or Δ*vhs* infection were excluded from further analysis.

### Processing of next-generation sequencing data

Sequencing reads were mapped against (i) the human genome (GRCh37/hg19), (ii) human rRNA sequences and (iii) the HSV-1 genome (HSV-1 strain 17, GenBank accession code: JN555585) using ContextMap v2.7.9 [64] (using BWA as short read aligner [65] and allowing a maximum indel size of 3 and at most 5 mismatches). For the two repeat regions in the HSV-1 genome, only one copy each was retained, excluding nucleotides 1–9,213 and 145,590–152,222. ContextMap produces unique mappings for each read, thus no further filtering was performed. Read coverage was visualized using Gviz [66] after normalizing to the total number of mapped human reads and averaging between replicates. For identification of enriched H3K4me3 regions (=peaks), BAM files with mapped reads were converted to BED format using BEDTools [67] (v2.24.0) and peaks were determined from BED files using F-Seq with default parameters [68]. Only peaks with length ≥500nt were considered. Unique non-overlapping peaks were identified by merging overlapping peaks across all samples using BEDTools. Overlaps of identified peaks to gene promoters were determined using ChIPseeker [69].

### Analysis of transcription read-through and differential gene expression

Number of read fragments per gene were determined from the mapped 4sU-seq and RNA-seq reads in a strand-specific manner using featureCounts [70] and gene annotations from Ensembl (version 87 for GRCh37/hg19) [71]. All fragments (read pairs for paired-end sequencing or reads for single-end sequencing) overlapping exonic regions on the corresponding strand by ≥25bp were counted for the corresponding gene. Expression of protein-coding genes and lincRNAs was quantified in terms of fragments per kilobase of exons per million mapped fragments (FPKM) and averaged between replicates. Only fragments mapping to the human genome were counted for the number of mapped fragments as previously described [19]. Downstream and upstream transcription for genes was determined from 4sU-seq data as described [20], i.e. the FPKM in the 5kb windows down- or upstream of genes divided by the gene FPKM. Read-through transcription was quantified as the difference in downstream transcription between infected and uninfected cells, with negative values set to zero. Read-in transcription was calculated analogously as the difference in upstream transcription between infected and uninfected cells. For full details, see our previous publication [20]. Only genes were included in this paper that (i) had no upstream or downstream gene within 5kb, (ii) were expressed (FPKM ≥ 1 in 4sU-RNA) in uninfected cells or at least one time point of WT infection and (iii) had at most 10% read-in transcription at any time during WT infection. For genes not expressed in uninfected cells (FPKM <1 in uninfected 4sU-RNA), at most 5% read-in transcription during infection and at most 25% upstream transcription in uninfected cells was allowed. These restrictions were used to exclude genes that only appeared induced due to read-in transcription from an upstream gene. In total, 4,162 genes were included for the analyses in this manuscript. Differential gene expression analysis for these genes in total and 4sU-RNA and subcellular RNA fractions was performed based on gene read counts using DESeq2 [22] and p-values were adjusted for multiple testing using the method by Benjamini and Hochberg [63]. Additional candidate up-regulated genes with low or no expression in uninfected cells were determined using the following criteria: i) FPKM in uninfected 4sU- and total RNA ≤ 1; ii) FPKM in either 4sU-RNA or total RNA at any time of infection both ≥ 0.5 and ≥4-fold higher than in uninfected cells; iii) read-in transcription ≤ 20% at all time points. Candidate genes were subsequently validated by manual inspection of mapped reads for individual replicates in the IGV genome browser [72]. To identify the extended set of *vhs*-dependently transcriptionally down-regulated genes, we applied DESeq2 for all genes on RNA-seq of chromatin-associated RNA in mock, 8h p.i WT and *Δvhs* infection. Genes were defined as transcriptionally down-regulated in a *vhs*-dependent manner if they were significantly down-regulated in WT (log2 fold-change ≤ −1, adj. p-value ≤ 0.001), not down-regulated in *Δvhs* infection (log2 fold-change > −1) and there was at least a 2-fold increase in fold-changes in *Δvhs* compared to WT infection.

### RNA half-lives

RNA half-lives were determined from total and 4sU-RNA FPKM values in uninfected cells as described [24] assuming a median RNA half-life of 5h.

### Mathematical model

The mathematical model of WT and Δ*vhs* infection is described in S1 Text.

### Clustering, enrichment and network analysis

Hierarchical clustering was performed in R [73] using Euclidean distances and Ward’s clustering criterion [74]. Gene Ontology (GO) [25] annotations for genes were obtained from EnrichR [75] and lists of interferon I, II and III up- or down-regulated genes (at least 2-fold) were obtained from the INTERFEROME database [26]. Genes regulated by doxycycline-inducible DUX4 were taken from the study of Jagannathan *et al*. (Supplementary Table 1; up-regulated: log2 fold-change ≥ 1, false discovery rate (fdr) ≤ 0.001; down-regulated: log2 fold-change ≤ −1, fdr ≤ 0.001) [49]. TCF-dependent genes and SRF targets in MEFs were taken from the study by Gualdrini *et al*. [40]. Odds-ratios and significance of enrichment compared to the background of 4,162 genes was determined using Fisher’s exact test in R [73] and p-values were adjusted for multiple testing using the method by Benjamini and Hochberg [63]. Human protein-protein associations were downloaded from the STRING database [35] (version 10.5) using NDEx [76] and visualized in Cytoscape [77]. Only associations with a score ≥350 are shown.

### Comparison of muSOX and vhs-dependent genes

Fold-changes for WT and ΔHS MHV68 infection were taken from the study of Abernathy *et al*. [17] and downloaded from Gene Expression Omnibus (GSE70481). Mouse and human gene symbols were mapped to their orthologs in the respective other species using the Mouse/Human Orthology table from the Mouse Genome Informatics (MGI) database [78]. muSOX-dependent genes were defined according to the criteria applied by Abernathy *et al*.: down-regulated in WT (log2 fold-change ≤ −1 and fdr ≤ 0.1) but not in ΔHS infection (log2 fold-change > −1 or fdr > 0.1). *vhs*-dependent genes were defined according to our criteria described above.

### Transcription factor binding motif search

Promoter motif search for *vhs*-dependently down-regulated genes was performed using HOMER in proximal promoter regions (−2,000 to +2,000 bp relative to the transcription start site). [79]. Potential transcription binding factor sites in uninfected cells were furthermore identified using ATAC-seq (Assay for Transposase-Accessible Chromatin using sequencing [80]) data of uninfected cells from our previous study (n=2 replicates) [20]. ATAC-seq data were mapped against hg19 as previously described [20] and open chromatin peaks were determined using MACS2 [81]. Blacklisted regions for hg19 (accession ENCFF001TDO) were downloaded from ENCODE [82] and peaks called in regions overlapping with blacklisted regions were removed from further analysis. Furthermore, only peaks occurring in both replicates were considered for motif search. Motif search was then performed using HOMER for open chromatin peaks within 10, 25 and 50kb, respectively, of *vhs*-dependently down-regulated genes.

## Supporting information captions

**S1 Text: Mathematical model**

Description and results of the mathematical model on *vhs* activity and loss of transcriptional activity in HSV-1 infection

**S2 File: Supplementary Figures**

Contains Supplementary Figures A-W and legends

**S3 Dataset: Read-through (in %) in WT and Δ*vhs* infection for genes included in the analysis**

**S4 Dataset: log2 fold-changes in all conditions for *vhs*-dependently transcriptionally down-regulated genes**

**S5 Dataset: Results of the GO term enrichment analysis for vhs-dependently transcriptionally downregulated genes**

p-values and odds-ratio were determined using Fisher’s exact test. p-values were adjusted for multiple testing using the method by Benjamini and Hochberg (BH). overlap = number of *vhs*-dependently downregulated genes annotated with GO term; not in list = number of *vhs*-dependently downregulated genes not annotated with GO term; background overlap = number of remaining genes annotated with GO term; background not in list = number of remaining genes not annotated with GO term; genes in overlap = *vhs*-dependently downregulated genes annotated with GO term.

**S6 Dataset: Extended set of *vhs*-dependently transcriptionally downregulated genes**

Log2 fold-changes and adjusted p-values in chromatin-associated RNA in 8h p.i. WT and Δvhs infection for the extended set of genes identified to be transcriptionally downregulated in a *vhs*-dependent manner.

**S7 Dataset: log2 fold-changes in all conditions and clusters for up-regulated genes**

**S8 Dataset: Unique non-overlapping H3K4me3 peak regions**

**S9 Dataset: Results of quantitative whole cell proteomics at 8h p.i. WT and Δvhs infection**

**S10 Dataset: Results of the GO term enrichment analysis for proteins down-regulated in both WT and *Δvhs* infection**

p-values and odds-ratio were determined using Fisher’s exact test. p-values were adjusted for multiple testing using the method by Benjamini and Hochberg (BH). overlap = number of down-regulated proteins annotated with GO term; not in list = number of down-regulated proteins not annotated with GO term; background overlap = number of remaining proteins annotated with GO term; background not in list = number of remaining proteins not annotated with GO term; genes in overlap = genes encoding for down-regulated proteins annotated with GO term.

**S11 Text: Full description of H3K4me3 ChIPmentation**

## References

1. Roizman B, Knipe DM, R.J. W. Herpes simplex viruses. In Knipe D M, Howley P M (ed), Fields virology, 5th ed Lippincott Williams & Wilkins, Philadelphia, PA. 2007:2501–601.

2. Kennedy PGE, Chaudhuri A. Herpes simplex encephalitis. Journal of Neurology, Neurosurgery & Psychiatry. 2002;73(3):237–8. doi: 10.1136/jnnp.73.3.237.

3. Kwong AD, Frenkel N. Herpes simplex virus-infected cells contain a function(s) that destabilizes both host and viral mRNAs. Proceedings of the National Academy of Sciences of the United States of America. 1987;84(7):1926–30. Epub 1987/04/01. PubMed PMID: 3031658; PubMed Central PMCID: PMC304554.

4. Oroskar AA, Read GS. Control of mRNA stability by the virion host shutoff function of herpes simplex virus. Journal of virology. 1989;63(5):1897–906. Epub 1989/05/01. PubMed PMID: 2539493; PubMed Central PMCID: PMC250601.

5. Feng P, Everly DN, Jr., Read GS. mRNA decay during herpesvirus infections: interaction between a putative viral nuclease and a cellular translation factor. Journal of virology. 2001;75(21):10272–80. Epub 2001/10/03. doi: 10.1128/JVI.75.21.10272-10280.2001. PubMed PMID: 11581395; PubMed Central PMCID: PMC114601.

6. Doepker RC, Hsu WL, Saffran HA, Smiley JR. Herpes simplex virus virion host shutoff protein is stimulated by translation initiation factors eIF4B and eIF4H. Journal of virology. 2004;78(9):4684–99. Epub 2004/04/14. PubMed PMID: 15078951; PubMed Central PMCID: PMC387725.

7. Sarma N, Agarwal D, Shiflett LA, Read GS. Small interfering RNAs that deplete the cellular translation factor eIF4H impede mRNA degradation by the virion host shutoff protein of herpes simplex virus. J Virol. 2008;82(13):6600–9. Epub 2008/05/02. doi: 10.1128/JVI.00137-08. PubMed PMID: 18448541; PubMed Central PMCID: PMC2447072.

8. Page HG, Read GS. The virion host shutoff endonuclease (UL41) of herpes simplex virus interacts with the cellular cap-binding complex eIF4F. Journal of virology. 2010;84(13):6886–90. Epub 2010/04/30. doi: 10.1128/JVI.00166-10JVI.00166-10 [pii]. PubMed PMID: 20427534; PubMed Central PMCID: PMC2903273.

9. Lam Q, Smibert CA, Koop KE, Lavery C, Capone JP, Weinheimer SP, et al. Herpes simplex virus VP16 rescues viral mRNA from destruction by the virion host shutoff function. EMBO J. 1996;15(10):2575–81. Epub 1996/05/15. PubMed PMID: 8665865; PubMed Central PMCID: PMC450190.

10. Taddeo B, Sciortino MT, Zhang W, Roizman B. Interaction of herpes simplex virus RNase with VP16 and VP22 is required for the accumulation of the protein but not for accumulation of mRNA. Proceedings of the National Academy of Sciences of the United States of America. 2007;104(29):12163–8. Epub 2007/07/11. doi: 0705245104 [pii] 10.1073/pnas.0705245104. PubMed PMID: 17620619; PubMed Central PMCID: PMC1924560.

11. Mbong EF, Woodley L, Dunkerley E, Schrimpf JE, Morrison LA, Duffy C. Deletion of the herpes simplex virus 1 UL49 gene results in mRNA and protein translation defects that are complemented by secondary mutations in UL41. J Virol. 2012;86(22):12351–61. Epub 2012/09/07. doi: 10.1128/JVI.01975-12. PubMed PMID: 22951838; PubMed Central PMCID: PMC3486455.

12. Shu M, Taddeo B, Zhang W, Roizman B. Selective degradation of mRNAs by the HSV host shutoff RNase is regulated by the UL47 tegument protein. Proc Natl Acad Sci U S A. 2013;110(18):E1669–75. Epub 2013/04/17. doi: 10.1073/pnas.1305475110. PubMed PMID: 23589852; PubMed Central PMCID: PMC3645526.

13. Spencer CA, Dahmus ME, Rice SA. Repression of host RNA polymerase II transcription by herpes simplex virus type 1. Journal of virology. 1997;71(3):2031–40. Epub 1997/03/01. PubMed PMID: 9032335; PubMed Central PMCID: PMC191289.

14. Abrisch RG, Eidem TM, Yakovchuk P, Kugel JF, Goodrich JA. Infection by Herpes Simplex Virus 1 Causes Near-Complete Loss of RNA Polymerase II Occupancy on the Host Cell Genome. Journal of virology. 2015;90(5):2503–13. doi: 10.1128/JVI.02665-15. PubMed PMID: 26676778; PubMed Central PMCID: PMCPMC4810688.

15. Birkenheuer CH, Danko CG, Baines JD. Herpes Simplex Virus 1 Dramatically Alters Loading and Positioning of RNA Polymerase II on Host Genes Early in Infection. J Virol. 2018. Epub 2018/02/14. doi: 10.1128/JVI.02184-17. PubMed PMID: 29437966; PubMed Central PMCID: PMC5874419.

16. Dai-Ju JQ, Li L, Johnson LA, Sandri-Goldin RM. ICP27 interacts with the C-terminal domain of RNA polymerase II and facilitates its recruitment to herpes simplex virus 1 transcription sites, where it undergoes proteasomal degradation during infection. J Virol. 2006;80(7):3567–81. Epub 2006/03/16. doi: 80/7/3567 [pii] 10.1128/JVI.80.7.3567-3581.2006. PubMed PMID: 16537625; PubMed Central PMCID: PMC1440381.

17. Abernathy E, Gilbertson S, Alla R, Glaunsinger B. Viral Nucleases Induce an mRNA Degradation-Transcription Feedback Loop in Mammalian Cells. Cell host & microbe. 2015;18(2):243–53. doi: https://doi.org/10.1016/j.chom.2015.06.019.

18. Pheasant K, Moller-Levet CS, Jones J, Depledge D, Breuer J, Elliott G. Nuclear-cytoplasmic compartmentalization of the herpes simplex virus 1 infected cell transcriptome is co-ordinated by the viral endoribonuclease vhs and cofactors to facilitate the translation of late proteins. PLoS Pathog. 2018;14(11):e1007331. Epub 2018/11/27. doi: 10.1371/journal.ppat.1007331. PubMed PMID: 30475899; PubMed Central PMCID: PMC6283614.

19. Rutkowski AJ, Erhard F, L’Hernault A, Bonfert T, Schilhabel M, Crump C, et al. Widespread disruption of host transcription termination in HSV-1 infection. Nature communications. 2015;6:7126. doi: 10.1038/ncomms8126. PubMed PMID: 25989971; PubMed Central PMCID: PMC4441252.

20. Hennig T, Michalski M, Rutkowski AJ, Djakovic L, Whisnant AW, Friedl MS, et al. HSV-1-induced disruption of transcription termination resembles a cellular stress response but selectively increases chromatin accessibility downstream of genes. PLoS Pathog. 2018;14(3):e1006954. Epub 2018/03/27. doi: 10.1371/journal.ppat.1006954. PubMed PMID: 29579120; PubMed Central PMCID: PMC5886697.

21. Whisnant AW, Jürges CS, Hennig T, Wyler E, Prusty B, Rutkowski AJ, et al. Integrative functional genomics decodes herpes simplex virus 1. Nature communications. 2020;11(1):2038. doi: 10.1038/s41467-020-15992-5.

22. Love MI, Huber W, Anders S. Moderated estimation of fold change and dispersion for RNA-seq data with DESeq2. Genome Biol. 2014;15(12):550. doi: 10.1186/s13059-014-0550-8. PubMed PMID: 25516281; PubMed Central PMCID: PMCPMC4302049.

23. Taddeo B, Esclatine A, Roizman B. The patterns of accumulation of cellular RNAs in cells infected with a wild-type and a mutant herpes simplex virus 1 lacking the virion host shutoff gene. Proceedings of the National Academy of Sciences. 2002;99(26):17031–6. doi: 10.1073/pnas.252588599.

24. Dölken L, Ruzsics Z, Radle B, Friedel CC, Zimmer R, Mages J, et al. High-resolution gene expression profiling for simultaneous kinetic parameter analysis of RNA synthesis and decay. RNA. 2008;14(9):1959–72.

25. The Gene Ontology Consortium. The Gene Ontology Resource: 20 years and still GOing strong. Nucleic acids research. 2019;47(D1):D330–D8. Epub 2018/11/06. doi: 10.1093/nar/gky1055. PubMed PMID: 30395331; PubMed Central PMCID: PMC6323945.

26. Rusinova I, Forster S, Yu S, Kannan A, Masse M, Cumming H, et al. Interferome v2.0: an updated database of annotated interferon-regulated genes. Nucleic acids research. 2013;41(Database issue):D1040–6. Epub 2012/12/04. doi: 10.1093/nar/gks1215. PubMed PMID: 23203888; PubMed Central PMCID: PMC3531205.

27. Sastry SK, Burridge K. Focal adhesions: a nexus for intracellular signaling and cytoskeletal dynamics. Experimental cell research. 2000;261(1):25–36. Epub 2000/11/18. doi: 10.1006/excr.2000.5043. PubMed PMID: 11082272.

28. Wang N, Butler JP, Ingber DE. Mechanotransduction across the cell surface and through the cytoskeleton. Science. 1993;260(5111):1124–7. Epub 1993/05/21. PubMed PMID: 7684161.

29. Horton ER, Byron A, Askari JA, Ng DHJ, Millon-Fremillon A, Robertson J, et al. Definition of a consensus integrin adhesome and its dynamics during adhesion complex assembly and disassembly. Nature cell biology. 2015;17(12):1577–87. Epub 2015/10/20. doi: 10.1038/ncb3257. PubMed PMID: 26479319; PubMed Central PMCID: PMC4663675.

30. Humphries JD, Byron A, Bass MD, Craig SE, Pinney JW, Knight D, et al. Proteomic analysis of integrin-associated complexes identifies RCC2 as a dual regulator of Rac1 and Arf6. Science signaling. 2009;2(87):ra51. Epub 2009/09/10. doi: 10.1126/scisignal.2000396. PubMed PMID: 19738201; PubMed Central PMCID: PMC2857963.

31. Robertson J, Jacquemet G, Byron A, Jones MC, Warwood S, Selley JN, et al. Defining the phospho-adhesome through the phosphoproteomic analysis of integrin signalling. Nature communications. 2015;6:6265. Epub 2015/02/14. doi: 10.1038/ncomms7265. PubMed PMID: 25677187; PubMed Central PMCID: PMC4338609.

32. Ng DH, Humphries JD, Byron A, Millon-Fremillon A, Humphries MJ. Microtubule-dependent modulation of adhesion complex composition. PLoS One. 2014;9(12):e115213. Epub 2014/12/20. doi: 10.1371/journal.pone.0115213. PubMed PMID: 25526367; PubMed Central PMCID: PMC4272306.

33. Schiller HB, Friedel CC, Boulegue C, Fassler R. Quantitative proteomics of the integrin adhesome show a myosin II-dependent recruitment of LIM domain proteins. EMBO reports. 2011;12(3):259–66. Epub 2011/02/12. doi: 10.1038/embor.2011.5. PubMed PMID: 21311561; PubMed Central PMCID: PMC3059911.

34. Schiller HB, Hermann MR, Polleux J, Vignaud T, Zanivan S, Friedel CC, et al. beta1- and alphav-class integrins cooperate to regulate myosin II during rigidity sensing of fibronectin-based microenvironments. Nature cell biology. 2013;15(6):625–36. Epub 2013/05/28. doi: 10.1038/ncb2747. PubMed PMID: 23708002.

35. Szklarczyk D, Morris JH, Cook H, Kuhn M, Wyder S, Simonovic M, et al. The STRING database in 2017: quality-controlled protein-protein association networks, made broadly accessible. Nucleic acids research. 2017;45(D1):D362–D8. Epub 2016/12/08. doi: 10.1093/nar/gkw937. PubMed PMID: 27924014; PubMed Central PMCID: PMC5210637.

36. Sarma N, Agarwal D, Shiflett LA, Read GS. Small interfering RNAs that deplete the cellular translation factor eIF4H impede mRNA degradation by the virion host shutoff protein of herpes simplex virus. Journal of virology. 2008;82(13):6600–9. Epub 2008/04/30. doi: 10.1128/JVI.00137-08. PubMed PMID: 18448541.

37. Fenwick ML, Everett RD. Inactivation of the Shutoff Gene (UL41) of Herpes Simplex Virus Types 1 and 2. Journal of General Virology. 1990;71(12):2961–7. doi: https://doi.org/10.1099/0022-1317-71-12-2961.

38. Buchwalter G, Gross C, Wasylyk B. Ets ternary complex transcription factors. Gene. 2004;324:1–14. Epub 2003/12/25. PubMed PMID: 14693367.

39. Schratt G, Philippar U, Berger J, Schwarz H, Heidenreich O, Nordheim A. Serum response factor is crucial for actin cytoskeletal organization and focal adhesion assembly in embryonic stem cells. J Cell Biol. 2002;156(4):737–50. Epub 2002/02/13. doi: 10.1083/jcb.200106008. PubMed PMID: 11839767; PubMed Central PMCID: PMC2174087.

40. Gualdrini F, Esnault C, Horswell S, Stewart A, Matthews N, Treisman R. SRF Co-factors Control the Balance between Cell Proliferation and Contractility. Mol Cell. 2016;64(6):1048–61. Epub 2016/11/22. doi: 10.1016/j.molcel.2016.10.016. PubMed PMID: 27867007; PubMed Central PMCID: PMC5179500.

41. Kamakura M, Goshima F, Luo C, Kimura H, Nishiyama Y. Herpes simplex virus induces the marked up-regulation of the zinc finger transcriptional factor INSM1, which modulates the expression and localization of the immediate early protein ICP0. Virology journal. 2011;8:257. Epub 2011/05/26. doi: 10.1186/1743-422X-8-257. PubMed PMID: 21609490; PubMed Central PMCID: PMC3125357.

42. Miyazaki D, Haruki T, Takeda S, Sasaki S, Yakura K, Terasaka Y, et al. Herpes simplex virus type 1-induced transcriptional networks of corneal endothelial cells indicate antigen presentation function. Investigative ophthalmology & visual science. 2011;52(7):4282–93. Epub 2011/05/05. doi: 10.1167/iovs.10-6911. PubMed PMID: 21540477.

43. Cheung P, Panning B, Smiley JR. Herpes simplex virus immediate-early proteins ICP0 and ICP4 activate the endogenous human alpha-globin gene in nonerythroid cells. Journal of virology. 1997;71(3):1784–93. Epub 1997/03/01. PubMed PMID: 9032307; PubMed Central PMCID: PMC191247.

44. Higgs DR, Hill AV, Bowden DK, Weatherall DJ, Clegg JB. Independent recombination events between the duplicated human alpha globin genes; implications for their concerted evolution. Nucleic acids research. 1984;12(18):6965–77. Epub 1984/09/25. PubMed PMID: 6091047; PubMed Central PMCID: PMC320136.

45. Schmidl C, Rendeiro AF, Sheffield NC, Bock C. ChIPmentation: fast, robust, low-input ChIP-seq for histones and transcription factors. Nat Methods. 2015;12(10):963–5. doi: 10.1038/nmeth.3542. PubMed PMID: 26280331; PubMed Central PMCID: PMCPMC4589892.

46. Lauberth SM, Nakayama T, Wu X, Ferris AL, Tang Z, Hughes SH, et al. H3K4me3 interactions with TAF3 regulate preinitiation complex assembly and selective gene activation. Cell. 2013;152(5):1021–36. Epub 2013/03/05. doi: 10.1016/j.cell.2013.01.052. PubMed PMID: 23452851; PubMed Central PMCID: PMC3588593.

47. Bernstein BE, Mikkelsen TS, Xie X, Kamal M, Huebert DJ, Cuff J, et al. A bivalent chromatin structure marks key developmental genes in embryonic stem cells. Cell. 2006;125(2):315–26. Epub 2006/04/25. doi: 10.1016/j.cell.2006.02.041. PubMed PMID: 16630819.

48. Full F, van Gent M, Sparrer KMJ, Chiang C, Zurenski MA, Scherer M, et al. Centrosomal protein TRIM43 restricts herpesvirus infection by regulating nuclear lamina integrity. Nature Microbiology. 2019;4(1):164–76. doi: 10.1038/s41564-018-0285-5.

49. Jagannathan S, Shadle SC, Resnick R, Snider L, Tawil RN, van der Maarel SM, et al. Model systems of DUX4 expression recapitulate the transcriptional profile of FSHD cells. Human Molecular Genetics. 2016;25(20):4419–31. doi: 10.1093/hmg/ddw271 %J Human Molecular Genetics.

50. Schwanhäusser B, Busse D, Li N, Dittmar G, Schuchhardt J, Wolf J, et al. Global quantification of mammalian gene expression control. Nature. 2011;473(7347):337–42. doi: 10.1038/nature10098.

51. Rodriguez W, Srivastav K, Muller M. C19ORF66 Broadly Escapes Virus-Induced Endonuclease Cleavage and Restricts Kaposi’s Sarcoma-Associated Herpesvirus. Journal of Virology. 2019;93(12):e00373–19. doi: 10.1128/jvi.00373-19.

52. Hu B, Li X, Huo Y, Yu Y, Zhang Q, Chen G, et al. Cellular responses to HSV-1 infection are linked to specific types of alterations in the host transcriptome. Sci Rep. 2016;6:28075. Epub 2016/06/30. doi: 10.1038/srep28075. PubMed PMID: 27354008; PubMed Central PMCID: PMC4926211.

53. Wyler E, Franke V, Menegatti J, Kocks C, Boltengagen A, Praktiknjo S, et al. Single-cell RNA-sequencing of herpes simplex virus 1-infected cells connects NRF2 activation to an antiviral program. Nature communications. 2019;10(1):4878. doi: 10.1038/s41467-019-12894-z.

54. Kamakura M, Nawa A, Ushijima Y, Goshima F, Kawaguchi Y, Kikkawa F, et al. Microarray analysis of transcriptional responses to infection by herpes simplex virus types 1 and 2 and their US3-deficient mutants. Microbes and infection. 2008;10(4):405–13. Epub 2008/04/12. doi: 10.1016/j.micinf.2007.12.019. PubMed PMID: 18403238.

55. Ziaie Z, Friedman HM, Kefalides NA. Suppression of matrix protein synthesis by herpes simplex virus type 1 in human endothelial cells. Collagen and related research. 1986;6(4):333–49. Epub 1986/10/01. PubMed PMID: 3028708.

56. Alpantaki K, Zafiropoulos A, Tseliou M, Vasarmidi E, Sourvinos G. Herpes simplex virus type-1 infection affects the expression of extracellular matrix components in human nucleus pulposus cells. Virus research. 2018;259:10–7. Epub 2018/10/20. doi: 10.1016/j.virusres.2018.10.010. PubMed PMID: 30339788.

57. Becker Y, Tavor E, Asher Y, Berkowitz C, Moyal M. Effect of herpes simplex virus type-1 UL41 gene on the stability of mRNA from the cellular genes: beta-actin, fibronectin, glucose transporter-1, and docking protein, and on virus intraperitoneal pathogenicity to newborn mice. Virus genes. 1993;7(2):133–43. Epub 1993/06/01. PubMed PMID: 8396282.

58. Morita T, Mayanagi T, Sobue K. Reorganization of the actin cytoskeleton via transcriptional regulation of cytoskeletal/focal adhesion genes by myocardin-related transcription factors (MRTFs/MAL/MKLs). Experimental cell research. 2007;313(16):3432–45. Epub 2007/08/24. doi: 10.1016/j.yexcr.2007.07.008. PubMed PMID: 17714703.

59. Sharili AS, Kenny FN, Vartiainen MK, Connelly JT. Nuclear actin modulates cell motility via transcriptional regulation of adhesive and cytoskeletal genes. Sci Rep. 2016;6:33893. Epub 2016/09/22. doi: 10.1038/srep33893. PubMed PMID: 27650314; PubMed Central PMCID: PMC5030641.

60. Fenwick ML, Everett RD. Inactivation of the shutoff gene (UL41) of herpes simplex virus types 1 and 2. J Gen Virol. 1990;71 (Pt 12):2961–7. Epub 1990/12/01. PubMed PMID: 2177088.

61. Tischer BK, Smith GA, Osterrieder N. En Passant Mutagenesis: A Two Step Markerless Red Recombination System. In: Braman J, editor. In Vitro Mutagenesis Protocols: Third Edition. Totowa, NJ: Humana Press; 2010. p. 421–30.

62. Sandbaumhüter M, Döhner K, Schipke J, Binz A, Pohlmann A, Sodeik B, et al. Cytosolic herpes simplex virus capsids not only require binding inner tegument protein pUL36 but also pUL37 for active transport prior to secondary envelopment. Cellular Microbiology. 2013;15(2):248–69. doi: 10.1111/cmi.12075.

63. Benjamini Y, Hochberg Y. Controlling the false discovery rate: a practical and powerful approach to multiple testing. Journal of the Royal Statistical Society, Series B. 1995;57(1):289–300.

64. Bonfert T, Kirner E, Csaba G, Zimmer R, Friedel CC. ContextMap 2: fast and accurate context-based RNA-seq mapping. BMC Bioinformatics. 2015;16:122. doi: 10.1186/s12859-015-0557-5. PubMed PMID: 25928589; PubMed Central PMCID: PMCPMC4411664.

65. Li H, Durbin R. Fast and accurate short read alignment with Burrows-Wheeler transform. Bioinformatics. 2009;25(14):1754–60. Epub 2009/05/20. doi: 10.1093/bioinformatics/btp324btp324 [pii]. PubMed PMID: 19451168; PubMed Central PMCID: PMC2705234.

66. Hahne F, Ivanek R. Visualizing Genomic Data Using Gviz and Bioconductor. In: Mathé E, Davis S, editors. Statistical Genomics: Methods and Protocols. New York, NY: Springer New York; 2016. p. 335–51.

67. Quinlan AR, Hall IM. BEDTools: a flexible suite of utilities for comparing genomic features. Bioinformatics. 2010;26(6):841–2. doi: 10.1093/bioinformatics/btq033. PubMed PMID: 20110278; PubMed Central PMCID: PMCPMC2832824.

68. Boyle AP, Guinney J, Crawford GE, Furey TS. F-Seq: a feature density estimator for high-throughput sequence tags. Bioinformatics. 2008;24(21):2537–8. doi: 10.1093/bioinformatics/btn480. PubMed PMID: 18784119; PubMed Central PMCID: PMCPMC2732284.

69. Yu G, Wang LG, He QY. ChIPseeker: an R/Bioconductor package for ChIP peak annotation, comparison and visualization. Bioinformatics. 2015;31(14):2382–3. doi: 10.1093/bioinformatics/btv145. PubMed PMID: 25765347.

70. Liao Y, Smyth GK, Shi W. featureCounts: an efficient general purpose program for assigning sequence reads to genomic features. Bioinformatics. 2014;30(7):923–30. doi: 10.1093/bioinformatics/btt656. PubMed PMID: 24227677.

71. Zerbino DR, Achuthan P, Akanni W, Amode MR, Barrell D, Bhai J, et al. Ensembl 2018. Nucleic acids research. 2018;46(D1):D754–D61. Epub 2017/11/21. doi: 10.1093/nar/gkx1098. PubMed PMID: 29155950; PubMed Central PMCID: PMC5753206.

72. Robinson JT, Thorvaldsdóttir H, Winckler W, Guttman M, Lander ES, Getz G, et al. Integrative genomics viewer. Nature biotechnology. 2011;29:24. doi: 10.1038/nbt.1754.

73. R Core Team. R: A Language and Environment for Statistical Computing. Vienna, Austria: R Foundation for Statistical Computing; 2018.

74. Murtagh F, Legendre P. Ward’s Hierarchical Agglomerative Clustering Method: Which Algorithms Implement Ward’s Criterion? Journal of Classification. 2014;31(3):274–95. doi: 10.1007/s00357-014-9161-z.

75. Chen EY, Tan CM, Kou Y, Duan Q, Wang Z, Meirelles GV, et al. Enrichr: interactive and collaborative HTML5 gene list enrichment analysis tool. BMC bioinformatics. 2013;14:128. Epub 2013/04/17. doi: 10.1186/1471-2105-14-128. PubMed PMID: 23586463; PubMed Central PMCID: PMC3637064.

76. Pratt D, Chen J, Welker D, Rivas R, Pillich R, Rynkov V, et al. NDEx, the Network Data Exchange. Cell systems. 2015;1(4):302–5. Epub 2015/11/26. doi: 10.1016/j.cels.2015.10.001. PubMed PMID: 26594663; PubMed Central PMCID: PMC4649937.

77. Shannon P, Markiel A, Ozier O, Baliga NS, Wang JT, Ramage D, et al. Cytoscape: a software environment for integrated models of biomolecular interaction networks. Genome Res. 2003;13(11):2498–504. Epub 2003/11/05. doi: 10.1101/gr.1239303. PubMed PMID: 14597658; PubMed Central PMCID: PMC403769.

78. Bult CJ, Blake JA, Smith CL, Kadin JA, Richardson JE, Group tMGD. Mouse Genome Database (MGD) 2019. Nucleic acids research. 2018;47(D1):D801–D6. doi: 10.1093/nar/gky1056.

79. Heinz S, Benner C, Spann N, Bertolino E, Lin YC, Laslo P, et al. Simple combinations of lineage-determining transcription factors prime cis-regulatory elements required for macrophage and B cell identities. Mol Cell. 2010;38(4):576–89. Epub 2010/06/02. doi: 10.1016/j.molcel.2010.05.004. PubMed PMID: 20513432; PubMed Central PMCID: PMC2898526.

80. Buenrostro JD, Wu B, Chang HY, Greenleaf WJ. ATAC-seq: A Method for Assaying Chromatin Accessibility Genome-Wide. Curr Protoc Mol Biol. 2015;109:21.9.1–.9.9. doi: 10.1002/0471142727.mb2129s109. PubMed PMID: 25559105.

81. Zhang Y, Liu T, Meyer CA, Eeckhoute J, Johnson DS, Bernstein BE, et al. Model-based analysis of ChIP-Seq (MACS). Genome biology. 2008;9(9):R137–R. Epub 2008/09/17. doi: 10.1186/gb-2008-9-9-r137. PubMed PMID: 18798982.

82. Encode Project Consortium. An integrated encyclopedia of DNA elements in the human genome. Nature. 2012;489(7414):57–74. Epub 2012/09/08. doi: 10.1038/nature11247. PubMed PMID: 22955616; PubMed Central PMCID: PMCPMC3439153.

